# Shared and distinct neural activity during anticipation and outcome of win and loss: A meta-analysis of the monetary incentive delay task

**DOI:** 10.1101/2022.07.21.500974

**Authors:** Yu Chen, Shefali Chaudhary, Chiang-Shan R. Li

## Abstract

Reward and punishment motivate decision making and behavioral changes. Numerous studies have examined regional activities during anticipation and outcome of win and loss in the monetary incentive delay task (MIDT). However, the great majority of studies reported findings of anticipation or outcome and of win or loss alone. It remains unclear how the neural correlates share and differentiate amongst these processes. We conducted an Activation Likelihood Estimation meta-analysis of 77 studies of the MIDT (5,779 subjects), including 24 published since the most recent meta-analysis, to identify and, with conjunction and subtraction, contrast regional responses to win anticipation, loss anticipation, win outcome, and loss outcome. Win and loss anticipation engaged a shared network of bilateral anterior insula (AI), striatum, thalamus, supplementary motor area (SMA), and precentral gyrus. Win and loss outcomes did not share regional activities. Win and loss outcome each engaged higher activity in medial orbitofrontal cortex (mOFC) and dorsal anterior cingulate cortex. Bilateral striatum and right occipital cortex responded to both anticipation and outcome of win, and right AI to both phases of loss. Win anticipation vs. outcome engaged higher activity in bilateral AI, striatum, SMA and precentral gyrus and right thalamus, and lower activity in bilateral mOFC and posterior cingulate cortex as well as right inferior frontal and angular gyri. Loss anticipation relative to outcome involved higher activity in bilateral striatum and left AI. These findings collectively suggest shared and distinct regional responses during monetary wins and losses. Delineating the neural correlates of these component processes may facilitate empirical research of motivated behavior and dysfunctional approach and avoidance in psychopathology.

**Highlights:**

- Win and loss anticipation both engaged the fronto-striatal-thalamic network.
- Win and loss outcomes shared no regional activities.
- The mOFC and dACC play distinct roles each in processing win and loss outcome.
- Win and loss anticipation engaged bilateral AI; loss outcome only the right AI.
- Win/loss anticipation vs. outcome engaged predominantly right/left AI.

## 1 Introduction

Reward- and punishment-driven decision making is fundamental to adaptive behaviors (Jean-Richard-Dit-Bressel et al., 2018; O’Doherty et al., 2017). Dysfunctional reward seeking and/or punishment avoidance have been implicated in many neuropsychiatric disorders (Whitton et al., 2015). For instance, depression patients showed lower sensitivity to rewards, impaired reward learning, and higher sensitivity to negative feedbacks (Admon and Pizzagalli, 2015; Eshel and Roiser, 2010). In contrast, antisocial personality disorder is characterized by elevated reward seeking and blunted punishment avoidance (Raine, 2018). Investigators have developed a variety of behavioral paradigms, including passive exposure to valenced stimuli, instrumental learning and reward-related decision-making (Richards et al., 2013), as well as the monetary incentive delay task (MIDT), to understand reward and punishment processing in health and illness (Balodis and Potenza, 2015; Knutson et al., 2001).

The MIDT allows investigations to distinguish between win and loss as well as between anticipation and consummation/outcome in reward processing. For example, the striatum and thalamus showed higher activation during anticipation of win vs. neutral outcomes (Dhingra et al., 2020; Dhingra et al., 2021; Knutson et al., 2001). The medial orbitofrontal cortex (mOFC), on the other hand, showed higher responses to win vs. neutral outcomes (Knutson et al., 2001; Treadway et al., 2013). Anticipation of loss vs. neutral outcomes engaged the ventral striatum, lateral thalamus, supplementary motor cortex, and insula (Bjork et al., 2010; Wu et al., 2014). Loss vs. neutral outcomes involved higher activations in the insula, inferior, middle, and superior frontal gyri, and superior parietal lobule (Maresh et al., 2014; Murray et al., 2020). Thus, previous studies have suggested potentially shared and distinct responses to the anticipation and outcome of wins and losses in the MIDT. Systematic reviews and meta-analyses of the studies can help in identifying the shared and distinct correlates.

A few meta-analyses of the MIDT have been published to investigate regional activations associated with win and loss processing. These meta-analyses have largely focused on the anticipation phase, likely because the great majority of fMRI studies reported solely the peak coordinates of win and loss anticipation in whole-brain analyses. An earlier meta-analysis demonstrated higher activation in the nucleus accumbens during win relative to loss anticipation and in the anterior insula during both win and loss (vs. nil) anticipation (Knutson and Greer, 2008). A more recent meta-analysis of 35 whole-brain (445 subjects) and 13 region of interest (254 subjects) studies highlighted shared response to loss anticipation and outcome (Dugre et al., 2018). Specifically, bilateral striatum, anterior insula, anterior cingulate cortex (ACC), and amygdala showed higher likelihood of activation during both loss anticipation and outcome. Oldham et al. (2018) identified regional responses to anticipation of wins (49 studies; 1,082 participants) and losses (32 studies; 681 participants), as well as to outcome of wins (22 studies; 691 participants). Specifically, the striatum, insula, amygdala, and thalamus showed higher activation when participants anticipated wins or losses (vs. nil), and the mOFC were recruited only during winning outcomes. Notably, none of the meta-analyses have systematically distinguished the shared and distinct correlates of valence and processing stage, namely win anticipation, loss anticipation, win outcome, and loss outcome. Distinguishing win and loss processing is clearly instrumental as the regional activities dictate opposing actions. Distinguishing anticipation and outcome phases of regional activities is also critical, with each reflecting the propensity to act and feedback about the action. These distinct component processes are fundamental to psychological models of adaptive learning.

To address this gap in research, we took advantage of a total of 24 additional studies published since the most recent and comprehensive meta-analysis of the MIDT (Oldham et al., 2018). We performed Activation Likelihood Estimation (ALE) to investigate the shared and distinct neural correlates underlying anticipation and outcome phases of win and loss processing and employed conjunction and subtraction analyses to identify regional activities that may overlap or differ between events of different valences and/or processing stages.

## 2 Methods

### 2.1 Literature search

Following the guidelines of “Preferred reporting items for systematic reviews and meta-analyses (PRISMA)”, we searched the literature on PubMed for imaging studies of MIDT with the key words “Monetary Incentive Delay Task” and “fMRI” and “NOT Review” and “NOT Meta-analysis”. We identified 338 studies on March 8, 2022. We also searched on Google Scholar and PsycNet (https://psycnet.apa.org/) using the same key words but found no new studies. A flow-chart for the procedure to arrive at the final sample for meta-analysis is shown in **Figure 1**. Only non-duplicate articles in English language (n = 330) were chosen for data mining if they included the following contrasts: anticipation of win vs. neutral (“win anticipation” hereafter), anticipation of loss vs. neutral (“loss anticipation”), win vs. neutral outcome (“win outcome”), or loss vs. neutral outcome (“loss outcome”). Patient studies were included if they contained data of healthy individuals. Likewise, medication or behavioral treatment studies were included if data of pre-treatment scans in healthy controls were available.

**Figure 1.**
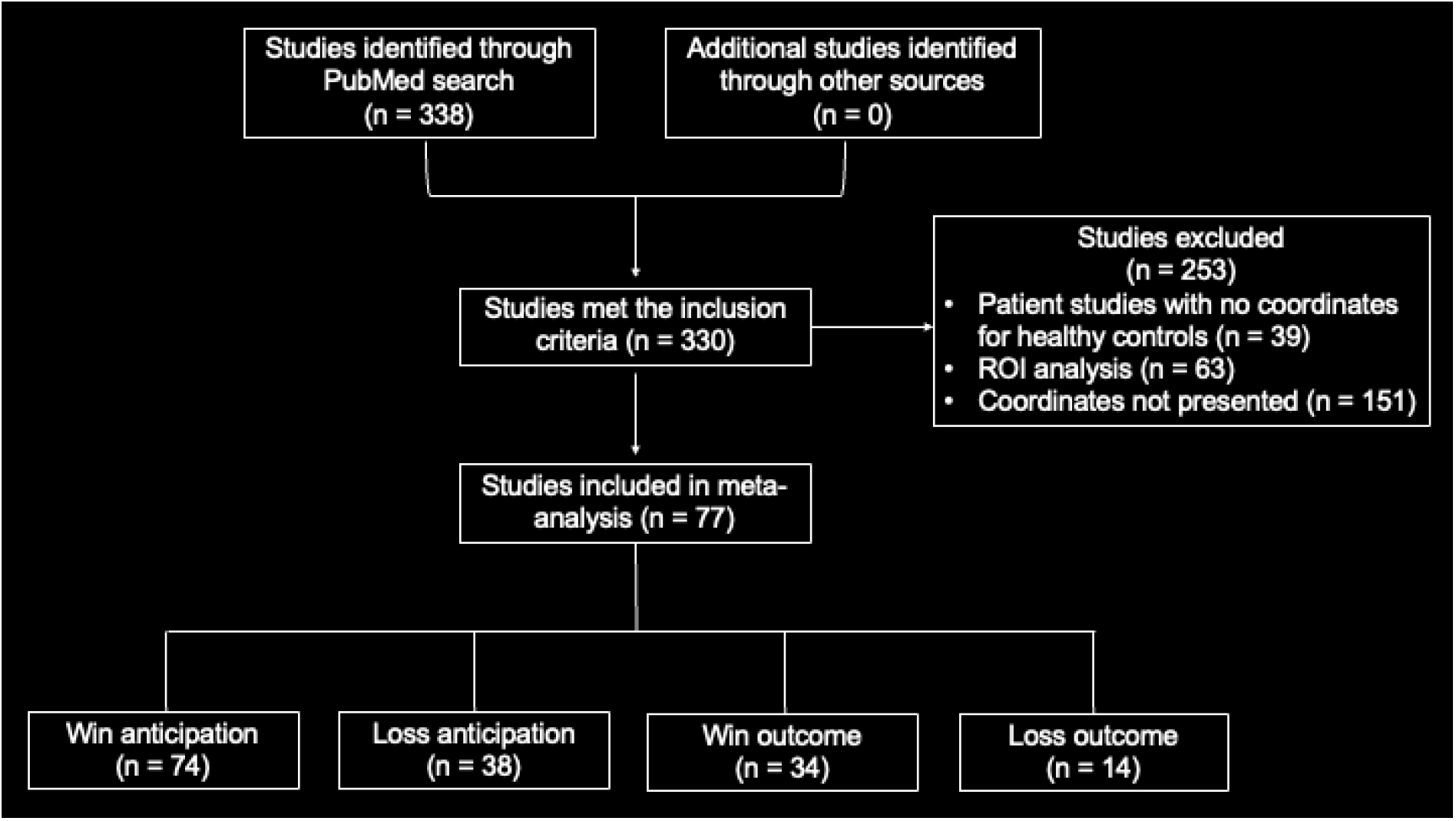
A flow-chart for the procedure to arrive at the final sample for meta-analysis, following ‘Preferred reporting items for systematic reviews and meta-analyses (PRISMA).’

Studies (n = 39) were removed based on the exclusion criteria, including life-time diagnosis of schizophrenia, depressive disorder, bipolar or manic disorder, or psychotic episodes, obsessive-compulsive disorder, post-traumatic stress disorder; treatment for mental disorders in the past 12 months, use of psychotropic medication; history of or current neurological disorders, major medical conditions, substance use, or brain trauma. Studies (n = 63) that used regional of interest (ROI) analyses were also excluded, which violated the assumption of ALE algorithm that each voxel in the entire brain has equal chance of being activated/showing correlation (Muller et al., 2018). One hundred and fifty-one studies were excluded because the coordinates of any of the four contrasts were not reported. If different rewards (e.g., $1 and $5) were used, the coordinates for the contrasts with the rewards combined (if available) or with the highest reward were used. A final pool of 77 studies of healthy volunteers were included in the current meta-analysis. A complete list of the studies is shown in **Supplementary Table S1**. Among the 77 studies, 74, 38, 34, and 14 reported peak coordinates (foci) of win anticipation, loss anticipation, win outcome, and loss outcome, respectively. We converted all foci that were reported in Talairach space to MNI space using the Lancaster transformation (Lancaster et al., 2007).

### 2.2 Activation likelihood estimation (ALE)

We used the GingerALE software package (version 3.0.2, http://brainmap.org/ale/) to perform ALE meta-analyses on coordinates in MNI space (Eickhoff et al., 2012; Eickhoff et al., 2009; Turkeltaub et al., 2012). The non-additive algorithm was used to reduce the bias of any single experiment (Turkeltaub et al., 2012). The ALE meta-analysis followed four main steps: computation of ALE scores, establishing a null distribution for statistical testing, thresholding, and cluster statistics, as described in detail in the GingerALE Manual (http://brainmap.org/ale/manual.pdf).

We performed the ALE single dataset analysis of each contrast - win anticipation, loss anticipation, win outcome, and loss outcome, using a cluster-forming threshold of voxel-level *p* < 0.001, uncorrected. Briefly, the non-additive ALE method was used to create a modelled activation (MA) map for each experiment (Turkeltaub et al., 2012) and a statistical whole-brain map was produced by combining all MA maps, where each voxel has an ALE value indicating its probability of activation. The resulting supra-threshold clusters were compared to a null distribution of cluster sizes established by 1,000 permutations of the data, at an FWE-corrected threshold of *p* < 0.05. We also performed ALE conjunction and subtraction analyses each to identify regional activities shared between contrasts and distinct to individual contrasts. The conjunction was created using the voxel-wise minimum value of the input ALE images as calculated in the single dataset analysis and the results were evaluated with a cluster-forming threshold of *p* < 0.001 uncorrected and a cluster-level threshold of *p* < 0.05 FWE corrected. The subtraction analysis was performed by repeating the following procedure for 5,000 times (Eickhoff et al., 2011): 1) GingerALE created simulated data by pooling the foci datasets and randomly dividing them into two groups of the same size as the original data set; 2) an ALE score was calculated at each voxel for each group; and 3) the difference between ALE scores was computed. The ALE values were collated across 5,000 permutations to yield an empirical null distribution for statistical inference. A *p*-value was assigned to each voxel based on how many times the difference in the null distribution exceeded the actual group difference. We applied a threshold of *p* < 0.001 uncorrected with a minimum cluster size of 100 mm^3^ to identify differences between any two contrasts, with the *Z*-score indicating the size of the differences at each voxel.

### 2.3 Evaluation of publication bias

We performed a “Fail-Safe N (FSN)” analysis to evaluate potential publication bias (Acar et al., 2018). We used the R program to generate a list of null studies with no statistically significant activation, all with a number of peaks and sample size equal to one of the studies in the original meta-analysis. The coordinates of these peaks were randomly drawn from the mask used by the ALE algorithm. For each single dataset analysis (i.e., win anticipation, loss anticipation, win outcome, and loss outcome), we computed the minimum numbers of null studies required in the FSN analysis – 5*k*+10 with *k* denoting the number of studies included in the original meta-analysis (Rosenthal, 1979). Specifically, at least 380, 200, 180, and 80 null studies were required for win anticipation, loss anticipation, win outcome, and loss outcome, respectively. We combined the original and these null studies and repeated the ALE meta-analyses. If the ALE findings remain significant, it means that results are sufficiently robust and are supported by at least the desired minimum of contributing studies. If adding a minimum of null studies alters the significant results of original ALE analyses, this indicates that results may not be robust when bias due to missing (noise) studies in the meta-analysis is present.

## 3 Results

### 3.1 Single dataset analyses

The results of ALE analyses of individual contrasts are shown in **Figure 2**. We found higher activation likelihood during win anticipation in bilateral midbrain regions (including red nucleus and superior colliculus), middle frontal gyri (MFG), supplementary motor area (SMA), anterior insula (AI), precentral gyri, occipital cortex (OC), thalamus, amygdala, and striatum (**Figure 2A**). Loss anticipation showed greater activation likelihood in bilateral SMA, AI, precentral gyri, thalamus, and striatum, and right amygdala (**Figure 2B**). Win outcome revealed clusters in bilateral mOFC, rostral anterior cingulate cortex (rACC), posterior cingulate cortex (PCC), striatum, and OC, left superior frontal gyrus, and right inferior frontal gyrus (**Figure 2C**). Loss outcome showed a large cluster in the AI extending to lateral posterior OFC, and temporal pole in the right hemisphere, and bilateral superior colliculus and dorsal ACC (**Figure 2D**). The clusters are summarized in **Supplementary Table S2**.

**Figure 2.**
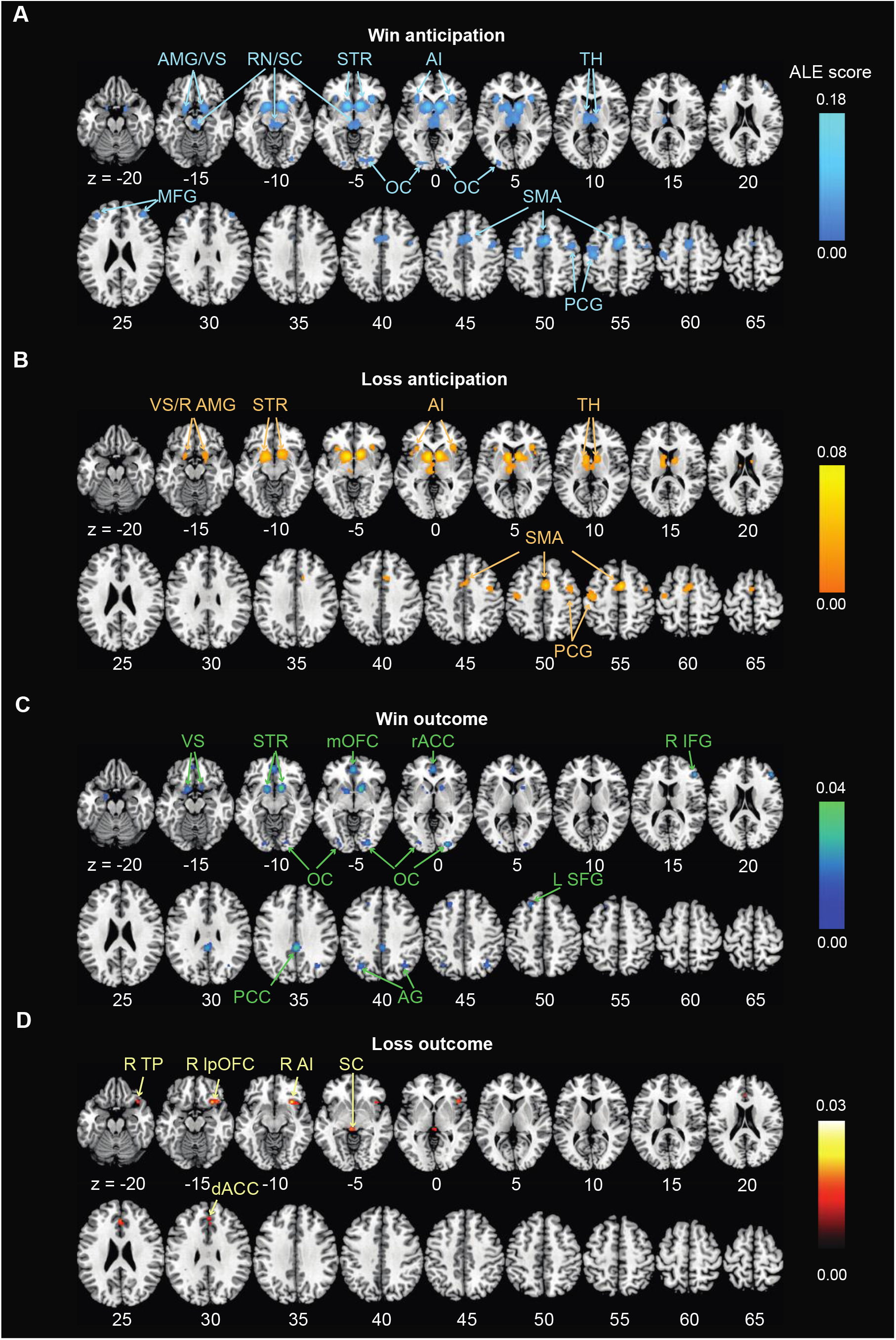
ALE single dataset analyses. **(A)** Win anticipation; **(B)** Loss anticipation; **(C)** Win outcome; and **(D)** Loss outcome. *Note:* The results were evaluated with a cluster-forming threshold of *p* < 0.001 uncorrected and a cluster-level threshold of *p* < 0.05 FWE corrected. Color bars represent ALE scores. L: left; R: right; ACC: anterior cingulate cortex; AI: anterior insula; AG: angular gyrus; AMG: amygdala; dACC: dorsal ACC; IFG: inferior frontal gyrus; lpOFC: lateral posterior orbitofrontal cortex; MFG: middle frontal gyrus; mOFC: medial orbitofrontal cortex; OC: occipital cortex; PCC: posterior cingulate cortex; PCG: precentral gyrus; rACC: rostral ACC; RN: red nucleus; SC: superior colliculus; SFG: superior frontal gyrus; SMA: supplementary motor area; STR: striatum; TH: thalamus; TP: temporal pole; VS: ventral striatum.

The publication bias was evaluated for individual contrasts, with additional null studies included in the ALE analyses. The ALE maps evaluated at the same threshold (**Supplementary Figure S1**) showed similar but fewer clusters in comparison with the original meta-analyses except that there were no significant findings for win outcome. The clusters are summarized in **Supplementary Table S5**. The findings indicated that the results for win anticipation, loss anticipation, and loss outcome were robust whereas the results for win outcome were subject to publication bias.

### 3.2 Conjunction and subtraction analyses

As shown in **Figure 3A**, win and loss anticipation shared activities in bilateral striatum, AI, precentral gyri, SMA, and thalamus. No clusters shared activities significantly between win and loss outcome. Win anticipation and outcome in conjunction showed clusters in bilateral striatum and right OC (**Figure 3C**). Loss anticipation and outcome in conjunction involved a small cluster of activity in the right AI extending to lateral OFC (**Figure 3D**). The clusters identified in conjunction analyses are summarized in **Supplementary Table S3**.

**Figure 3.**
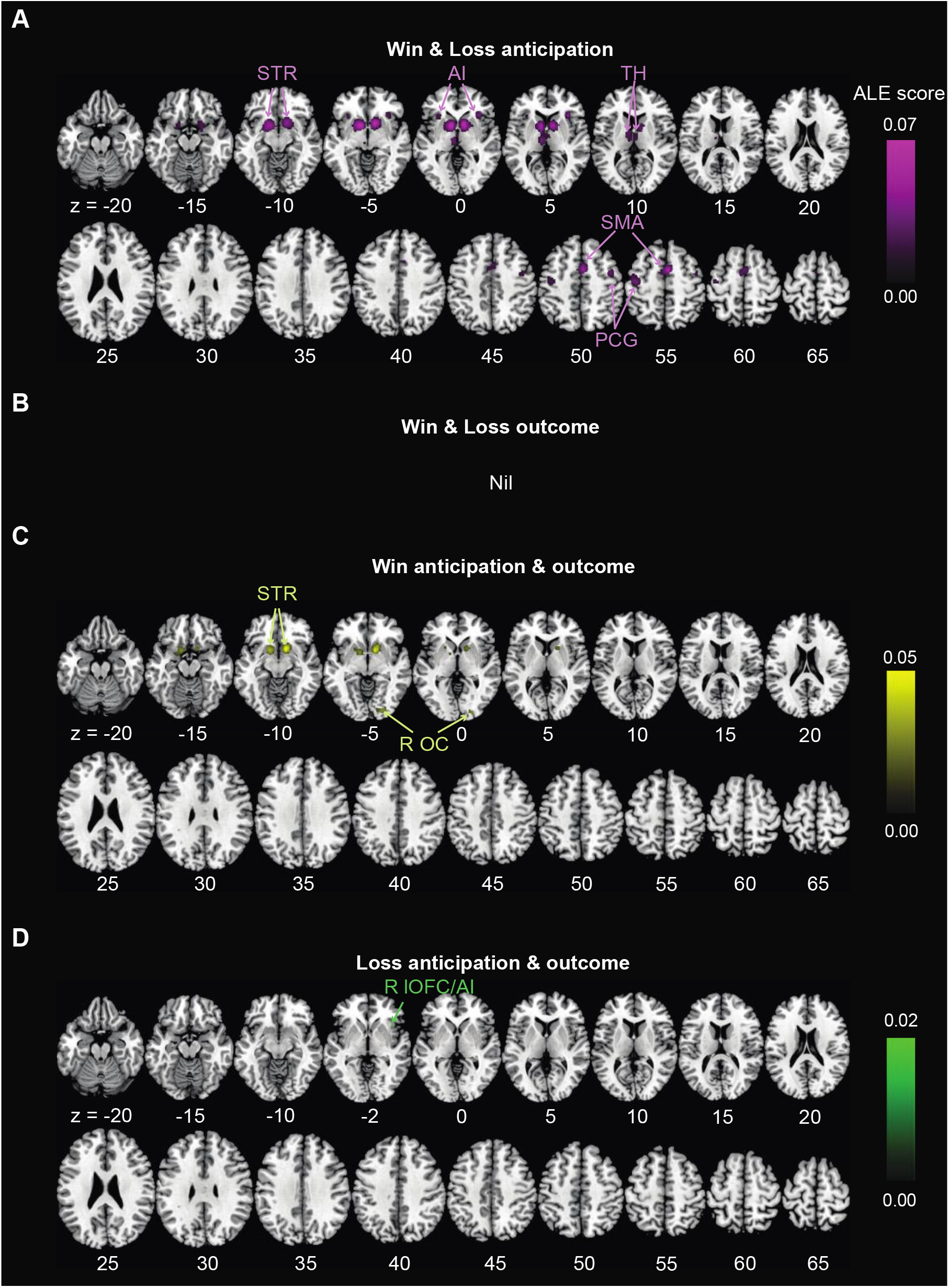
ALE conjunction analyses: **(A)** Win and Loss anticipation; **(B)** Win and Loss outcome; **(C)** Win anticipation and outcome; and **(D)** Loss anticipation and outcome. The results were evaluated with a cluster-forming threshold of *p* < 0.001 uncorrected and a cluster-level threshold of *p* < 0.05 FWE corrected. Color bars represent ALE scores. Nil: no significant findings. R: right; AI: anterior insula; OC: occipital cortex; PCG: precentral gyrus; SMA: supplementary motor area; STR: striatum; TH: thalamus.

As shown in **Figure 4**, win and loss anticipation showed no significant differences in activity. Win relative to loss outcome showed higher activation likelihood in bilateral mOFC (**Figure 4B**). Loss relative to win outcome revealed no significant differences. Win anticipation vs. outcome showed higher activity in bilateral AI (but predominantly right AI), striatum, SMA, and precentral gyri, and right thalamus. Win outcome vs. anticipation, on the contrary, activated bilateral mOFC and PCC, as well as right inferior frontal and angular gyri (**Figure 4C**). Loss anticipation relative to outcome involved bilateral striatum, and left AI, while loss outcome vs. anticipation showed no significant differences (**Figure 4D**). The clusters identified in subtraction analyses are summarized in **Supplementary Table S4**.

**Figure 4.**
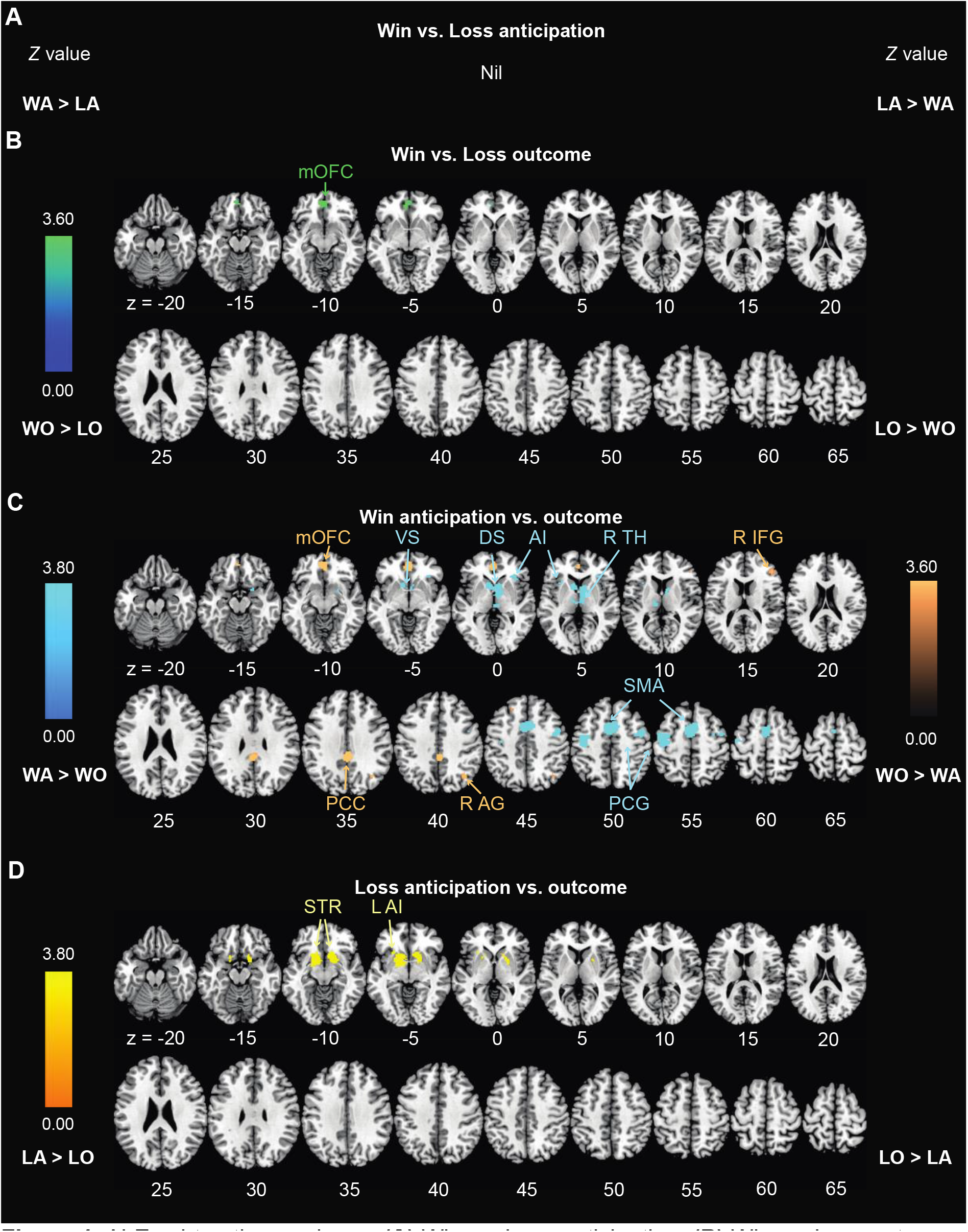
ALE subtraction analyses: **(A)** Win vs. Loss anticipation; **(B)** Win vs. Loss outcome; **(C)** Win anticipation vs. outcome; and **(D)** Loss anticipation vs. outcome. *Note:* Subtraction analyses were conducted with a significance level of *p* < 0.001 with a minimal cluster size of 100 mm^3^. Color bars represent *Z* values. Nil: no significant findings; WA: win anticipation; WO: win outcome; LA: loss anticipation; LO: loss outcome. L: left; R: right; AG: angular gyrus; AI: anterior insula; DS: dorsal striatum; IFG: inferior frontal gyrus; mOFC: medial orbitofrontal cortex; PCC: posterior cingulate cortex; PCG: precentral gyrus; SMA: supplementary motor area; STR: striatum; TH: thalamus; VS: ventral striatum.

## 4 Discussion

To our knowledge, this is the first sufficiently powered meta-analysis of whole-brain MIDT studies to investigate regional brain responses to the anticipatory and consummatory phases of win and loss processing. The processes engaged both shared and distinct neural correlates, in line with previous findings (Dugre et al., 2018; Knutson and Greer, 2008; Liu et al., 2011; Oldham et al., 2018). With a larger number of studies included for meta-analysis, we observed regional activities, particularly those related to loss outcome, that were not reported previously. Further, with conjunction and subtraction analyses, we identified regional activities that were significantly different between valences and across processing stages. Specifically, win and loss anticipation both engaged the fronto-striatal-thalamic networks; in contrast, win and loss outcomes shared no regional activities. The mOFC and dACC play specific roles each in processing win and loss outcome. Win anticipation and outcome both engaged bilateral VS and the right-hemispheric OC, whereas loss anticipation and outcome both involved higher activity in the right lateral OFC and AI. Win anticipation vs. outcome involved higher activity in their shared front-striatal-thalamic network and lower activity in mOFC, PCC, and right AG, regions of the default mode network. Notably, win anticipation vs. outcome involved higher activity in bilateral but predominantly right AI, whereas loss anticipation vs. outcome involved higher left AI activity. In the below, we provided an overview of the regional activities with reference to previous studies (Section 4.1) and discussed the shared (4.2) and distinct (4.3) correlates.

### 4.1 Neural correlates of win/loss anticipation and outcome

We replicated the findings for win and loss anticipation in the most recently published meta-analysis of MIDT (Oldham et al., 2018), showing activations across a wide swath of brain regions, including the amygdala, midbrain, striatum, AI, thalamus, SMA, precentral gyrus, and OC. Moreover, we observed activation of other frontal cortical regions, including the MFG for win anticipation. Encompassing behavioral tasks that included the MIDT, a recent meta-analysis also showed activation of MFG and SMA during win anticipation (Jauhar et al., 2021). Although the roles of the MFG in win/loss processing have not been investigated systematically, prior studies implicated the MFG in motivated behaviors (Bahlmann et al., 2015). The MFG was activated in reward (high > low) × cognitive load (high > low) interaction during goal-directed behaviors (Pochon et al., 2002; Taylor et al., 2004). Moreover, individual approach and avoidance traits were associated with activation of left- and right-lateralized MFG, respectively, during Stroop conflicts (Spielberg et al., 2011). Here, we observed bilateral MFG activation during win anticipation only; thus, the behavioral contexts that support functional lateralization of the MFG remains to be clarified.

We observed activation of the SMA for both win and loss anticipation, consistent with a role of this medial frontal region in behavioral selection based on stimulus-reward associations and representation of risky decisions involving a potential loss (Bickel et al., 2009; Hartstra et al., 2010). The SMA encodes reward expectancy, as demonstrated with neuronal recordings in monkeys (Campos et al., 2005; Lee, 2004). In a task that required physical effort (grip force) to receive money reward, a network comprising the SMA and caudal dACC showed higher activation to larger reward and to smaller effort. Further, higher SMA activation was correlated with choice of less effortful rather than higher reward options, suggesting that SMA biased choices on the bases of motor costs rather than reward (Klein-Flugge et al., 2016). Thus, the functions of SMA in motivated behavior may go beyond simply signaling the magnitude of reward at stake and involve more nuanced evaluation of cost and trade-off between effort and reward.

We observed that the rACC, inferior and superior frontal gyri, angular gyrus, and OC, in addition to the striatum, amygdala, mOFC, and PCC as demonstrated by Oldham et al. (2018), showed higher likelihood of activation for win vs. nil outcome. For loss vs. nil outcome, we found activation in the temporal pole, lateral posterior OFC, AI, dACC, and superior colliculus, predominantly in the right hemisphere. The latter findings contrasted with bilateral putamen and globus pallidum reported in Dugre et al. (2018), which included both whole-brain and ROI studies and employed effect-size seed-based *d* Mapping, instead of ALE, for meta-analysis. Notably, we identified mOFC and lateral OFC each during win and loss outcome, consistent with previous evidence that reward and punishment are represented medially and laterally, respectively, in the OFC during reversal learning (O’Doherty et al., 2001).

### 4.2 Shared neural correlates during win and loss processing

We found that both VS and dorsal striatum (DS) contribute substantially to win and loss anticipation as well as to win outcome, in accord with Oldham et al. (2018). Previous studies suggested functional heterogeneity within striatal subregions, with the VS involved in encoding both positive and negative stimuli and the DS in associative and motor aspects of decision-making (Burton et al., 2015). Studies in rats also showed elevated neuronal activities in the VS and DS each in association with the expectation of larger rewards and behavioral responses to retrieve the reward (Burton et al., 2014; Roesch et al., 2009). During Pavlovian conditioning, the VS was critical in learning motivationally salient stimuli, independent of valence, to bias action selection (Jensen et al., 2007). Therefore, the VS may encode salience during the anticipatory period and modulate motivational processes in the DS to initiate the pursuit of reward or to avoid loss (Burton et al., 2015; Oldham et al., 2018). The findings here and of previous studies that the striatum responds to anticipation of wins and losses may reflect the fact that avoiding monetary loss is equivalent to winning in the MIDT. Studies that distinguish reward and punishment (e.g., with electric shocks) categorically may be needed to differentiate striatal responses to anticipation of positive and negative outcomes.

Previous meta-analysis identified the VS and left amygdala as common correlates during win anticipation and outcome at a threshold of *p* < 0.005 (Oldham et al., 2018). Our conjunction analyses did not show the left amygdala at *p* < 0.001 but did at *p* < 0.005 (results not shown). Our findings further showed a higher likelihood of activation of right OC in the conjunction of win anticipation and outcome. In addition to processing visual information (Op de Beeck and Baker, 2010), the OC is involved in encoding emotional salience and motivation (Geday et al., 2003; Sabatinelli et al., 2011), as during reward conditioning (Kirsch et al., 2003) and passive exposure to pictures of food vs. objects (Schur et al., 2009). The OC also showed higher activation during decision-making under risky and uncertain but not certain conditions, suggesting its broad engagement in behavioral responses to saliency (Blankenstein et al., 2017; Guo et al., 2013).

With conjunction analysis, we showed shared responses of the right AI (rAI) to loss anticipation and outcome. The rAI showed stronger activation during risky vs. safe choices in decision making and its activity during risky choices was significantly correlated with the likelihood of selecting a safe response after punishment and with higher individual scores of harm avoidance (Paulus et al., 2003). An MIDT study reported rAI activity during anticipation of large (but not small) losses in association with individual traits of negative (but not positive) emotional arousal (Wu et al., 2014). These along with the current findings suggest an outsized role of the rAI in loss processing and behavioral avoidance.

### 4.3 Distinct neural correlates of win and loss processing

The mOFC play important roles in motivational and emotional regulation (Rempel-Clower, 2007; Rudebeck and Rich, 2018). Here, we demonstrated higher likelihood of activation of the mOFC in response to win but not loss outcome. The differences in regional activities were confirmed by the subtraction analysis of win > loss outcome, broadly in line with mOFC response to reward but not punishment across multiple behavioral tasks (O’Doherty et al., 2001; Rolls, 2019; Rolls et al., 2020). Subtraction analysis also showed that the mOFC was more likely to be activated during win outcome vs. anticipation, as reported earlier by Oldham et al (2018). This is consistent with the finding from behaving monkeys that mOFC neurons rapidly encoded the value of a selected action and continued to signal the outcome until after its delivery in a two-option gambling task (Strait et al., 2014). As discussed earlier, medial and lateral OFC respond to rewarding and punishing outcomes, respectively. Although the subtraction analysis failed to reveal lateral OFC activity in loss vs. win outcome, investigators should revisit this issue as studies of MIDT accrue in the literature.

We showed higher likelihood of activation of the dACC to loss outcome but not anticipation, although subtraction analysis did not substantiate the differences. The dACC has been implicated in integrating and learning the risk of an action to optimize decision-making (Bush et al., 2002; Kennerley et al., 2006; Rushworth et al., 2004). The dACC showed higher activation during decisions to quit vs. to chase losses (Campbell-Meiklejohn et al., 2008), suggesting dACC’s role in processing negative outcomes for behavioral adjustment. Prior studies also demonstrated co-activation of the rAI and dACC during the anticipation of an electric shock (Chua et al., 1999). More broadly, both rAI and dACC showed higher activation to social exclusion vs. inclusion in the Cyberball task (Moor et al., 2012). Here, with the rAI responding to both loss anticipation and outcome and the dACC only to loss outcome, future work may investigate how rAI and dACC dynamically interact in loss processing for behavioral control.

The thalamus showed higher likelihood of activation during the anticipatory period only, regardless of valence, although the phase specificity was confirmed for win but not loss with the subtraction analyses of anticipation vs. outcome. Oldham et al. (2018) too observed thalamic activity during win anticipation vs. outcome. Also in support were findings that rodent thalamic neurons elevated firing as reward values increased during the delay period, peaking before the delivery of reward, suggesting reward anticipation and prediction (Komura et al., 2001). These findings are nonetheless surprising given the role of the thalamus in processing and relaying sensory inputs to the cortex (Sherman and Guillery, 2006) and in salience detection (Matsumoto et al., 2001), irrespective of valence (Kirouac, 2015). Neurons in the thalamus of mice encoded the saliency of both appetitive or aversive outcomes, and the inhibition of these thalamic responses suppressed appetitive or aversive associative learning and extinction (Zhu et al., 2018). In dynamic causal modeling of the MIDT data, an earlier work proposed a functional circuit of incentive processing where anticipation of win or loss generated “alerting” signals in the thalamus that integrate with interoceptive information conveyed by the AI to shape action selection in the striatum (Cho et al., 2013). It was also proposed that the thalamus receives inputs from the striatum and in turn projects to the PFC, thereby linking reward signals to “higher-order” cognitive functions (Rademacher et al., 2010). In other studies, neuronal responses in the thalamus, global pallidus, and ACC were parametrically modulated by reward levels only whereas parametric responses to both reward and punishment were observed in bilateral insula, caudate head, and OFC (Elliott et al., 2000). Thus, the thalamus shows higher likelihood of activation during anticipation and the differences between anticipation and outcome activities are most evident during reward processing. Future research may address the effective connectivity within the thalamic-striatal-insular/frontal networks for both reward and loss processing, to better understand how the regional activities and connectivities support motivated behavior, including those involved in drug seeking (Li et al., in press; Naqvi and Bechara, 2009).

Win anticipation vs. outcome involved higher activity in their shared fronto-striatal-thalamic network and lower activity in mOFC, PCC, and right AG, consistent with opposing patterns of activity of the executive control and default mode networks (DMN; Raichle, 2015). These findings should also be considered along with DMN regional reactivity to motivationally salient stimuli (Breiter et al., 2001; Dohmatob et al., 2020; Mohanty et al., 2008; Pearson et al., 2011; Rogers et al., 2004). In monkeys, neurons in the PCC showed transient phasic increases in firing during the detection of salient environmental changes (Hayden et al., 2009), such as the delivery of a reward (McCoy et al., 2003). In humans, the PCC responds to motivational salience of the target in guiding shifts of spatial attention (Dohmatob et al., 2020). Indeed, here we also observed higher mOFC, PCC, and right AG activity during win outcome vs. nil (Figure 2), which appeared to drive the difference in these DMN regional activities between win anticipation and outcome.

A few findings suggest functional lateralization of the AI. First, whereas both win and loss anticipation engaged bilateral AI, loss outcome engaged the right AI only and win outcome engaged neither right nor left AI. As a result, win anticipation vs. outcome involved higher activity in bilateral but predominantly right AI, whereas loss anticipation vs. outcome involved higher left AI activity. In earlier reviews Craig and colleagues suggested that the right and left AI responds to aversive and to positive and affiliative emotions, respectively (Craig, 2009; Craig, 2005). For instance, in healthy individuals, pleasant vs. unpleasant music (Koelsch et al., 2006) as well as anticipation of (Simmons et al., 2004) and exposure to (Straube and Miltner, 2011) emotionally aversive vs. neutral pictures activated the right but not left AI. However, these findings contrast with other reports associating greater responses of the left AI with visually aversive vs. neutral stimuli and higher negative valence ratings of the stimuli (Caria et al., 2010) as well as meta-analysis linking predominantly right and left insula activity each to approach/positive emotion and withdrawal/negative emotion-related behaviors (Wager et al., 2003). The latter reports appeared to be consistent with our finding that win anticipation vs. outcome involved higher activity in bilateral but predominantly right AI, whereas loss anticipation vs. outcome involved higher left AI activity. Together, these observations along with studies showing sensitivity of the AI to uncertainty (Fan et al., 2014; Wu et al., 2021) suggest the potential importance in considering the processing phase of anticipation and consummation in elucidating functional lateralization of the AI.

### 4.4 Limitations of the study, other considerations, and conclusions

A few limitations should be acknowledged. Firstly, the studies reporting the contrasts of loss events are fewer in number than those reporting win events, which may have impacted the statistical power of ALE analyses. With more MIDT studies to investigate the neural mechanism of loss processing, meta-analyses are to follow up on the roles of shared and distinct regional response to wins and losses. Secondly, an earlier meta-analysis distinguished VS sensitivity to reward magnitudes during both prediction and consumption and mOFC sensitivity only during consumption (Diekhof et al., 2012). Very few MIDT studies reported coordinates for the contrasts of different magnitudes; thus, we were not able to verify these differences. Finally, a recent meta-analysis demonstrated that adolescents vs. adults showed higher likelihood for activation in the insula, striatum, amygdala, ACC, and OFC during reward processing (Silverman et al., 2015). As the reward circuits “mature” before self-regulation circuits during this developmental period (Casey, 2015; Casey et al., 2008; Chen et al., 2020), it would be of instrumental importance to investigate how win and loss processes and their neural mechanisms evolve from adolescence to adulthood.

In conclusion, we demonstrated both shared and non-shared neural correlates of anticipatory and consummatory win and loss processing. The findings highlighted that while win and loss outcomes shared no regional activities, win and loss anticipation both engaged the fronto-striatal-thalamic network; the mOFC and dACC play distinct roles each in processing win and loss outcome; and win anticipation vs. outcome engaged bilateral but predominantly right AI, whereas loss anticipation vs. outcome involved higher left AI activity.

## Acknowledgements

This study is supported by NIH grants AG067024 and AG072893. The funding agencies are otherwise not responsible for the design of the study, data collection or analysis, or in the decision to publish these results.

## Data/code availability statement

Data used in the meta-analysis were collected from literature and are provided in the Supplement. This research did not generate any codes.

## References

Acar, F., Seurinck, R., Eickhoff, S.B., Moerkerke, B., 2018. Assessing robustness against potential publication bias in Activation Likelihood Estimation (ALE) meta-analyses for fMRI. PLoS One 13, e0208177.

Admon, R., Pizzagalli, D.A., 2015. Dysfunctional Reward Processing in Depression. Curr Opin Psychol 4, 114–118.

Bahlmann, J., Aarts, E., D’Esposito, M., 2015. Influence of motivation on control hierarchy in the human frontal cortex. J Neurosci 35, 3207–3217.

Balodis, I.M., Potenza, M.N., 2015. Anticipatory reward processing in addicted populations: a focus on the monetary incentive delay task. Biol Psychiatry 77, 434–444.

Bickel, W.K., Pitcock, J.A., Yi, R., Angtuaco, E.J., 2009. Congruence of BOLD response across intertemporal choice conditions: fictive and real money gains and losses. J Neurosci 29, 8839–8846.

Bjork, J.M., Smith, A.R., Chen, G., Hommer, D.W., 2010. Adolescents, adults and rewards: comparing motivational neurocircuitry recruitment using fMRI. PLoS One 5, e11440.

Blankenstein, N.E., Peper, J.S., Crone, E.A., van Duijvenvoorde, A.C.K., 2017. Neural Mechanisms Underlying Risk and Ambiguity Attitudes. J Cogn Neurosci 29, 1845–1859.

Breiter, H.C., Aharon, I., Kahneman, D., Dale, A., Shizgal, P., 2001. Functional imaging of neural responses to expectancy and experience of monetary gains and losses. Neuron 30, 619–639.

Burton, A.C., Bissonette, G.B., Lichtenberg, N.T., Kashtelyan, V., Roesch, M.R., 2014. Ventral striatum lesions enhance stimulus and response encoding in dorsal striatum. Biol Psychiatry 75, 132–139.

Burton, A.C., Nakamura, K., Roesch, M.R., 2015. From ventral-medial to dorsal-lateral striatum: neural correlates of reward-guided decision-making. Neurobiol Learn Mem 117, 51–59.

Bush, G., Vogt, B.A., Holmes, J., Dale, A.M., Greve, D., Jenike, M.A., Rosen, B.R., 2002. Dorsal anterior cingulate cortex: a role in reward-based decision making. Proc Natl Acad Sci U S A 99, 523–528.

Campbell-Meiklejohn, D.K., Woolrich, M.W., Passingham, R.E., Rogers, R.D., 2008. Knowing when to stop: the brain mechanisms of chasing losses. Biol Psychiatry 63, 293–300.

Campos, M., Breznen, B., Bernheim, K., Andersen, R.A., 2005. Supplementary motor area encodes reward expectancy in eye-movement tasks. J Neurophysiol 94, 1325–1335.

Caria, A., Sitaram, R., Veit, R., Begliomini, C., Birbaumer, N., 2010. Volitional control of anterior insula activity modulates the response to aversive stimuli. A real-time functional magnetic resonance imaging study. Biol Psychiatry 68, 425–432.

Casey, B.J., 2015. Beyond simple models of self-control to circuit-based accounts of adolescent behavior. Annu Rev Psychol 66, 295–319.

Casey, B.J., Getz, S., Galvan, A., 2008. The adolescent brain. Dev Rev 28, 62–77.

Chen, Y., Chen, C., Wu, T., Qiu, B., Zhang, W., Fan, J., 2020. Accessing the development and heritability of the capacity of cognitive control. Neuropsychologia 139, 107361.

Cho, Y.T., Fromm, S., Guyer, A.E., Detloff, A., Pine, D.S., Fudge, J.L., Ernst, M., 2013. Nucleus accumbens, thalamus and insula connectivity during incentive anticipation in typical adults and adolescents. Neuroimage 66, 508–521.

Chua, P., Krams, M., Toni, I., Passingham, R., Dolan, R., 1999. A functional anatomy of anticipatory anxiety. Neuroimage 9, 563–571.

Craig, A., 2009. How do you feel--now? The anterior insula and human awareness. Nature reviews neuroscience 10.

Craig, A.D., 2005. Forebrain emotional asymmetry: a neuroanatomical basis? Trends Cogn Sci 9, 566–571.

Dhingra, I., Zhang, S., Zhornitsky, S., Le, T.M., Wang, W., Chao, H.H., Levy, I., Li, C.-S.R., 2020. The effects of age on reward magnitude processing in the monetary incentive delay task. Neuroimage 207, 116368.

Dhingra, I., Zhang, S., Zhornitsky, S., Wang, W., Le, T.M., Li, C.-S.R., 2021. Sex differences in neural responses to reward and the influences of individual reward and punishment sensitivity. BMC neuroscience 22, 1–14.

Diekhof, E.K., Kaps, L., Falkai, P., Gruber, O., 2012. The role of the human ventral striatum and the medial orbitofrontal cortex in the representation of reward magnitude - an activation likelihood estimation meta-analysis of neuroimaging studies of passive reward expectancy and outcome processing. Neuropsychologia 50, 1252–1266.

Dohmatob, E., Dumas, G., Bzdok, D., 2020. Dark control: The default mode network as a reinforcement learning agent. Hum Brain Mapp 41, 3318–3341.

Dugre, J.R., Dumais, A., Bitar, N., Potvin, S., 2018. Loss anticipation and outcome during the Monetary Incentive Delay Task: a neuroimaging systematic review and meta-analysis. PeerJ 6, e4749.

Eickhoff, S.B., Bzdok, D., Laird, A.R., Kurth, F., Fox, P.T., 2012. Activation likelihood estimation meta-analysis revisited. Neuroimage 59, 2349–2361.

Eickhoff, S.B., Bzdok, D., Laird, A.R., Roski, C., Caspers, S., Zilles, K., Fox, P.T., 2011. Co-activation patterns distinguish cortical modules, their connectivity and functional differentiation. Neuroimage 57, 938–949.

Eickhoff, S.B., Laird, A.R., Grefkes, C., Wang, L.E., Zilles, K., Fox, P.T., 2009. Coordinate-based activation likelihood estimation meta-analysis of neuroimaging data: a random-effects approach based on empirical estimates of spatial uncertainty. Hum Brain Mapp 30, 2907–2926.

Elliott, R., Friston, K.J., Dolan, R.J., 2000. Dissociable neural responses in human reward systems. Journal of Neuroscience 20, 6159–6165.

Eshel, N., Roiser, J.P., 2010. Reward and punishment processing in depression. Biological Psychiatry 68, 118–124.

Fan, J., Van Dam, N.T., Gu, X., Liu, X., Wang, H., Tang, C.Y., Hof, P.R., 2014. Quantitative characterization of functional anatomical contributions to cognitive control under uncertainty. Journal of cognitive neuroscience 26, 1490–1506.

Geday, J., Gjedde, A., Boldsen, A.-S., Kupers, R., 2003. Emotional valence modulates activity in the posterior fusiform gyrus and inferior medial prefrontal cortex in social perception. Neuroimage 18, 675–684.

Guo, Z., Chen, J., Liu, S., Li, Y., Sun, B., Gao, Z., 2013. Brain areas activated by uncertain reward-based decision-making in healthy volunteers. Neural Regen Res 8, 3344–3352.

Hartstra, E., Oldenburg, J.F., Van Leijenhorst, L., Rombouts, S.A., Crone, E.A., 2010. Brain regions involved in the learning and application of reward rules in a two-deck gambling task. Neuropsychologia 48, 1438–1446.

Hayden, B.Y., Smith, D.V., Platt, M.L., 2009. Electrophysiological correlates of default-mode processing in macaque posterior cingulate cortex. Proc Natl Acad Sci U S A 106, 5948–5953.

Jauhar, S., Fortea, L., Solanes, A., Albajes-Eizagirre, A., McKenna, P.J., Radua, J., 2021. Brain activations associated with anticipation and delivery of monetary reward: A systematic review and meta-analysis of fMRI studies. PLoS One 16, e0255292.

Jean-Richard-Dit-Bressel, P., Killcross, S., McNally, G.P., 2018. Behavioral and neurobiological mechanisms of punishment: implications for psychiatric disorders. Neuropsychopharmacology 43, 1639–1650.

Jensen, J., Smith, A.J., Willeit, M., Crawley, A.P., Mikulis, D.J., Vitcu, I., Kapur, S., 2007. Separate brain regions code for salience vs. valence during reward prediction in humans. Hum Brain Mapp 28, 294–302.

Kennerley, S.W., Walton, M.E., Behrens, T.E., Buckley, M.J., Rushworth, M.F., 2006. Optimal decision making and the anterior cingulate cortex. Nat Neurosci 9, 940–947.

Kirouac, G.J., 2015. Placing the paraventricular nucleus of the thalamus within the brain circuits that control behavior. Neurosci Biobehav Rev 56, 315–329.

Kirsch, P., Schienle, A., Stark, R., Sammer, G., Blecker, C., Walter, B., Ott, U., Burkart, J., Vaitl, D., 2003. Anticipation of reward in a nonaversive differential conditioning paradigm and the brain reward system. Neuroimage 20, 1086–1095.

Klein-Flugge, M.C., Kennerley, S.W., Friston, K., Bestmann, S., 2016. Neural Signatures of Value Comparison in Human Cingulate Cortex during Decisions Requiring an Effort-Reward Trade-off. J Neurosci 36, 10002–10015.

Knutson, B., Fong, G.W., Adams, C.M., Varner, J.L., Hommer, D., 2001. Dissociation of reward anticipation and outcome with event-related fMRI. Neuroreport 12, 3683–3687.

Knutson, B., Greer, S.M., 2008. Anticipatory affect: neural correlates and consequences for choice. Philos Trans R Soc Lond B Biol Sci 363, 3771–3786.

Koelsch, S., Fritz, T., Dy, V.C., Muller, K., Friederici, A.D., 2006. Investigating emotion with music: an fMRI study. Hum Brain Mapp 27, 239–250.

Komura, Y., Tamura, R., Uwano, T., Nishijo, H., Kaga, K., Ono, T., 2001. Retrospective and prospective coding for predicted reward in the sensory thalamus. Nature 412, 546–549.

Lancaster, J.L., Tordesillas-Gutierrez, D., Martinez, M., Salinas, F., Evans, A., Zilles, K., Mazziotta, J.C., Fox, P.T., 2007. Bias between MNI and Talairach coordinates analyzed using the ICBM-152 brain template. Hum Brain Mapp 28, 1194–1205.

Lee, D., 2004. Behavioral context and coherent oscillations in the supplementary motor area. J Neurosci 24, 4453–4459.

Li, G., Chen, Y., Tang, X., Li, C.R., in press. Loss and frontal striatal reactivities interrelate alcohol use severity and rule-breaking behavior in young adult drinkers. Biological psychiatry: cognitive neuroscience and neuroimaging.

Liu, X., Hairston, J., Schrier, M., Fan, J., 2011. Common and distinct networks underlying reward valence and processing stages: a meta-analysis of functional neuroimaging studies. Neurosci Biobehav Rev 35, 1219–1236.

Maresh, E.L., Allen, J.P., Coan, J.A., 2014. Increased default mode network activity in socially anxious individuals during reward processing. Biology of mood & anxiety disorders 4, 1–12.

Matsumoto, N., Minamimoto, T., Graybiel, A.M., Kimura, M., 2001. Neurons in the thalamic CM-Pf complex supply striatal neurons with information about behaviorally significant sensory events. Journal of neurophysiology 85, 960–976.

McCoy, A.N., Crowley, J.C., Haghighian, G., Dean, H.L., Platt, M.L., 2003. Saccade reward signals in posterior cingulate cortex. Neuron 40, 1031–1040.

Mohanty, A., Gitelman, D.R., Small, D.M., Mesulam, M.M., 2008. The spatial attention network interacts with limbic and monoaminergic systems to modulate motivation-induced attention shifts. Cereb Cortex 18, 2604–2613.

Moor, B.G., Guroglu, B., Op de Macks, Z.A., Rombouts, S.A., Van der Molen, M.W., Crone, E.A., 2012. Social exclusion and punishment of excluders: neural correlates and developmental trajectories. Neuroimage 59, 708–717.

Muller, V.I., Cieslik, E.C., Laird, A.R., Fox, P.T., Radua, J., Mataix-Cols, D., Tench, C.R., Yarkoni, T., Nichols, T.E., Turkeltaub, P.E., Wager, T.D., Eickhoff, S.B., 2018. Ten simple rules for neuroimaging meta-analysis. Neurosci Biobehav Rev 84, 151–161.

Murray, L., Lopez-Duran, N.L., Mitchell, C., Monk, C.S., Hyde, L.W., 2020. Neural mechanisms of reward and loss processing in a low-income sample of at-risk adolescents. Soc Cogn Affect Neurosci 15, 1310–1325.

Naqvi, N.H., Bechara, A., 2009. The hidden island of addiction: the insula. Trends Neurosci 32, 56–67.

O’Doherty, J., Kringelbach, M.L., Rolls, E.T., Hornak, J., Andrews, C., 2001. Abstract reward and punishment representations in the human orbitofrontal cortex. Nature neuroscience 4, 95–102.

O’Doherty, J.P., Cockburn, J., Pauli, W.M., 2017. Learning, Reward, and Decision Making. Annu Rev Psychol 68, 73–100.

Oldham, S., Murawski, C., Fornito, A., Youssef, G., Yucel, M., Lorenzetti, V., 2018. The anticipation and outcome phases of reward and loss processing: A neuroimaging meta-analysis of the monetary incentive delay task. Hum Brain Mapp 39, 3398–3418.

Op de Beeck, H.P., Baker, C.I., 2010. The neural basis of visual object learning. Trends Cogn Sci 14, 22–30.

Paulus, M.P., Rogalsky, C., Simmons, A., Feinstein, J.S., Stein, M.B., 2003. Increased activation in the right insula during risk-taking decision making is related to harm avoidance and neuroticism. Neuroimage 19, 1439–1448.

Pearson, J.M., Heilbronner, S.R., Barack, D.L., Hayden, B.Y., Platt, M.L., 2011. Posterior cingulate cortex: adapting behavior to a changing world. Trends Cogn Sci 15, 143–151.

Pochon, J.B., Levy, R., Fossati, P., Lehericy, S., Poline, J.B., Pillon, B., Le Bihan, D., Dubois, B., 2002. The neural system that bridges reward and cognition in humans: an fMRI study. Proc Natl Acad Sci U S A 99, 5669–5674.

Rademacher, L., Krach, S., Kohls, G., Irmak, A., Grunder, G., Spreckelmeyer, K.N., 2010. Dissociation of neural networks for anticipation and consumption of monetary and social rewards. Neuroimage 49, 3276–3285.

Raichle, M.E., 2015. The brain’s default mode network. Annu Rev Neurosci 38, 433–447.

Raine, A., 2018. Antisocial personality as a neurodevelopmental disorder. Annual Review of Clinical Psychology 14, 259–289.

Rempel-Clower, N.L., 2007. Role of orbitofrontal cortex connections in emotion. Ann N Y Acad Sci 1121, 72–86.

Richards, J.M., Plate, R.C., Ernst, M., 2013. A systematic review of fMRI reward paradigms used in studies of adolescents vs. adults: the impact of task design and implications for understanding neurodevelopment. Neurosci Biobehav Rev 37, 976–991.

Roesch, M.R., Singh, T., Brown, P.L., Mullins, S.E., Schoenbaum, G., 2009. Ventral striatal neurons encode the value of the chosen action in rats deciding between differently delayed or sized rewards. J Neurosci 29, 13365–13376.

Rogers, R.D., Ramnani, N., Mackay, C., Wilson, J.L., Jezzard, P., Carter, C.S., Smith, S.M., 2004. Distinct portions of anterior cingulate cortex and medial prefrontal cortex are activated by reward processing in separable phases of decision-making cognition. Biol Psychiatry 55, 594–602.

Rolls, E.T., 2019. The orbitofrontal cortex and emotion in health and disease, including depression. Neuropsychologia 128, 14–43.

Rolls, E.T., Cheng, W., Feng, J., 2020. The orbitofrontal cortex: reward, emotion and depression. Brain Commun 2, fcaa196.

Rosenthal, R., 1979. The file drawer problem and tolerance for null results. Psychological bulletin 86.

Rudebeck, P.H., Rich, E.L., 2018. Orbitofrontal cortex. Curr Biol 28, R1083–R1088.

Rushworth, M.F., Walton, M.E., Kennerley, S.W., Bannerman, D.M., 2004. Action sets and decisions in the medial frontal cortex. Trends Cogn Sci 8, 410–417.

Sabatinelli, D., Fortune, E.E., Li, Q., Siddiqui, A., Krafft, C., Oliver, W.T., Beck, S., Jeffries, J., 2011. Emotional perception: meta-analyses of face and natural scene processing. Neuroimage 54, 2524–2533.

Schur, E.A., Kleinhans, N.M., Goldberg, J., Buchwald, D., Schwartz, M.W., Maravilla, K., 2009. Activation in brain energy regulation and reward centers by food cues varies with choice of visual stimulus. Int J Obes (Lond) 33, 653–661.

Sherman, S.M., Guillery, R.W., 2006. Exploring the thalamus and its role in cortical function. MIT press.

Silverman, M.H., Jedd, K., Luciana, M., 2015. Neural networks involved in adolescent reward processing: An activation likelihood estimation meta-analysis of functional neuroimaging studies. Neuroimage 122, 427–439.

Simmons, A., Matthews, S.C., Stein, M.B., Paulus, M.P., 2004. Anticipation of emotionally aversive visual stimuli activates right insula. Neuroreport 15, 2261–2265.

Spielberg, J.M., Miller, G.A., Engels, A.S., Herrington, J.D., Sutton, B.P., Banich, M.T., Heller, W., 2011. Trait approach and avoidance motivation: lateralized neural activity associated with executive function. Neuroimage 54, 661–670.

Strait, C.E., Blanchard, T.C., Hayden, B.Y., 2014. Reward value comparison via mutual inhibition in ventromedial prefrontal cortex. Neuron 82, 1357–1366.

Straube, T., Miltner, W.H., 2011. Attention to aversive emotion and specific activation of the right insula and right somatosensory cortex. Neuroimage 54, 2534–2538.

Taylor, S.F., Welsh, R.C., Wager, T.D., Phan, K.L., Fitzgerald, K.D., Gehring, W.J., 2004. A functional neuroimaging study of motivation and executive function. Neuroimage 21, 1045–1054.

Treadway, M.T., Buckholtz, J.W., Zald, D.H., 2013. Perceived stress predicts altered reward and loss feedback processing in medial prefrontal cortex. Front Hum Neurosci 7, 180.

Turkeltaub, P.E., Eickhoff, S.B., Laird, A.R., Fox, M., Wiener, M., Fox, P., 2012. Minimizing within-experiment and within-group effects in activation likelihood estimation meta-analyses. Human brain mapping 33, 1–13.

Wager, T.D., Phan, K.L., Liberzon, I., Taylor, S.F., 2003. Valence, gender, and lateralization of functional brain anatomy in emotion: a meta-analysis of findings from neuroimaging. Neuroimage 19, 513–531.

Whitton, A.E., Treadway, M.T., Pizzagalli, D.A., 2015. Reward processing dysfunction in major depression, bipolar disorder and schizophrenia. Curr Opin Psychiatry 28, 7–12.

Wu, C.C., Samanez-Larkin, G.R., Katovich, K., Knutson, B., 2014. Affective traits link to reliable neural markers of incentive anticipation. Neuroimage 84, 279–289.

Wu, T., Schulz, K.P., Fan, J., 2021. Activation of the cognitive control network associated with information uncertainty. Neuroimage 230, 117703.

Zhu, Y., Nachtrab, G., Keyes, P.C., Allen, W.E., Luo, L., Chen, X., 2018. Dynamic salience processing in paraventricular thalamus gates associative learning. Science 362, 423–429.

