## Supplement for "Shared and distinct neural activity during anticipation and outcome of win and loss: A meta-analysis of the monetary incentive delay task"

**Supplementary Table S1.** A list of 77 studies included in the meta-analysis

| No. | Study | n | Win anticipation | Loss anticipation | Win outcome | Loss outcome |
|---|---|---|---|---|---|---|
| 1 | Abler et al. (2007) | 8 | Y | N | Y | N |
| 2 | Adcock et al. (2006) | 12 | Y | N | N | N |
| 3 | Asari et al. (2018) | 19 | Y | Y | N | N |
| 4 | Balodis et al. (2012) | 14 | Y | Y | Y | Y |
| 5 | Barker et al. (2021) | 874 | Y | N | N | N |
| 6 | Barman et al. (2015) | 63 | Y | N | N | N |
| 7 | Behan et al. (2015) | 20 | Y | N | N | N |
| 8 | Bhutani et al. (2021) | 76 | Y | N | Y | N |
| 9 | Bjork et al. (2010) | 48 | Y | Y | Y | Y |
| 10 | Bretzke et al. (2021) | 47 | Y | N | N | N |
| 11 | Cao et al. (2019) | 1510 | Y | N | N | N |
| 12 | Carl et al. (2016) | 20 | Y | N | Y | N |
| 13 | Carter et al. (2009) | 17 | Y | Y | N | N |
| 14 | Cho et al. (2013) | 50 | Y | Y | N | N |
| 15 | Cohn et al. (2015) | 128 | Y | Y | Y | Y |
| 16 | Colich et al. (2017) | 76 | Y | Y | N | N |
| 17 | Cope et al. (2019) | 34 | Y | Y | N | N |
| 18 | Costumero et al. (2013) | 44 | Y | N | N | N |
| 19 | Damiano et al. (2014) | 31 | Y | N | N | N |
| 20 | Dillon et al. (2010) | 31 | Y | N | Y | N |
| 21 | Dunlop et al. (2020) | 87 | Y | N | Y | N |
| 22 | Edel et al. (2013) | 12 | Y | N | N | N |
| 23 | Enzi et al. (2012) | 15 | Y | Y | N | N |
| 24 | Figee et al. (2011) | 19 | Y | N | Y | N |
| 25 | Filbey et al. (2013) | 27 | N | N | Y | Y |
| 26 | Funayama et al. (2014) | 20 | Y | Y | N | N |
| 27 | Gonzalez et al. (2016) | 83 | Y | Y | N | N |
| 28 | Grimm et al. (2021) | 45 | Y | N | N | N |
| 29 | Hanssen et al. (2015) | 57 | Y | N | Y | N |
| 30 | He et al. (2019) | 20 | Y | Y | Y | Y |
| 31 | Herbort et al. (2016) | 23 | Y | Y | N | N |
| 32 | Ikeda et al. (2019) | 15 | Y | Y | N | N |
| 33 | Johnson et al. (2019) | 24 | Y | Y | Y | N |
| 34 | Juckel et al. (2006) | 10 | Y | Y | N | N |
| 35 | Juckel et al. (2012) | 13 | Y | Y | N | N |
| 36 | Kappel et al. (2013) | 20 | Y | N | N | N |
| 37 | Kaufmann et al. (2013) | 19 | Y | Y | N | N |
| 38 | Kim et al. (2020) | 18 | Y | Y | N | N |
| 39 | Kirk et al. (2015) | 44 | Y | Y | Y | N |
| 40 | Knutson et al. (2008) | 12 | Y | Y | Y | Y |
| 41 | Kocsel et al. (2019) | 41 | Y | Y | Y | Y |
| 42 | Kocsel et al. (2017) | 37 | Y | Y | Y | Y |
| 43 | Kollmann et al. (2017) | 41 | Y | Y | N | N |

| | | | | | | |
|---|---|---|---|---|---|---|
| 44 | Li et al. (2019) | 174 | Y | Y | N | N |
| 45 | Maresh et al. (2014) | 84 | Y | Y | Y | Y |
| 46 | Martz et al. (2021) | 145 | Y | N | N | N |
| 47 | Martz et al. (2018) | 235 | Y | N | N | N |
| 48 | Metzak et al. (2021) | 179 | Y | N | N | N |
| 49 | Michielse et al. (2019) | 40 | Y | N | N | N |
| 50 | Montoya et al. (2014) | 20 | Y | N | N | N |
| 51 | Morelli et al. (2021) | 46 | Y | N | Y | N |
| 52 | Mucci et al. (2015) | 22 | Y | Y | Y | N |
| 53 | Murray et al. (2020) | 128 | Y | Y | Y | Y |
| 54 | Navas et al. (2018) | 68 | Y | N | Y | N |
| 55 | Nymberg et al. (2013) | 143 | Y | N | N | N |
| 56 | Ossewaarde et al. (2011) | 28 | Y | N | N | N |
| 57 | Paraskevopoulou et al. (2021) | 40 | Y | N | Y | N |
| 58 | Pfabigan et al. (2014) | 25 | Y | Y | N | N |
| 59 | Rademacher et al. (2010) | 28 | Y | N | Y | N |
| 60 | Saji et al. (2013) | 18 | Y | Y | N | N |
| 61 | Schlagenhauf et al. (2008) | 10 | Y | N | N | N |
| 62 | Simon et al. (2010) | 24 | Y | N | Y | N |
| 63 | Simon et al. (2015) | 27 | N | N | Y | N |
| 64 | Stoy et al. (2011) | 12 | Y | Y | N | N |
| 65 | Stoy et al. (2012) | 15 | Y | Y | N | N |
| 66 | Strohle et al. (2008) | 10 | Y | N | Y | N |
| 67 | Treadway et al. (2013) | 38 | Y | Y | Y | Y |
| 68 | van Hell et al. (2012) | 11 | Y | N | Y | N |
| 69 | Verdejo-Roman et al. (2017) | 76 | Y | N | Y | N |
| 70 | Veroude et al. (2016) | 103 | Y | N | Y | N |
| 71 | Weidacker et al. (2021) | 20 | N | Y | N | Y |
| 72 | Weiland et al. (2014) | 12 | Y | N | N | N |
| 73 | Wu et al. (2014) | 52 | Y | Y | Y | N |
| 74 | Yan et al. (2016) | 22 | Y | Y | Y | Y |
| 75 | Yao et al. (2020) | 30 | Y | Y | Y | Y |
| 76 | Yau et al. (2012) | 40 | Y | Y | N | N |
| 77 | Zweynert et al. (2011) | 30 | Y | N | N | N |

*Note:* Y: yes; N: no.

**Supplementary Table S2.** ALE results for activations during win anticipation, loss anticipation, win outcome, and loss outcome

| **Cluster volumes (mm$^3$)** | **ALE** | **MNI Coordinates (mm)** | | | **Identified Region** |
|---|---|---|---|---|---|
| | | **x** | **y** | **z** | |
| ***Win anticipation*** | | | | | |
| 33,464 | 0.178 | 10 | 10 | -2 | Right Caudate Head |
| | 0.146 | -10 | 8 | -4 | Left Caudate Head |
| | 0.060 | 4 | -12 | 6 | Right Thalamus, Medial Dorsal Nucleus |
| | 0.057 | -10 | -16 | 8 | Left Thalamus, Medial Dorsal Nucleus |
| | 0.048 | 6 | -26 | -6 | Right Midbrain, Red Nucleus |
| | 0.044 | -4 | -26 | -4 | Left Midbrain, Red Nucleus |
| | 0.034 | -2 | -18 | -14 | Left Midbrain, Red Nucleus |

| | | | | | |
|---|---|---|---|---|---|
| | 0.025 | -20 | -2 | 12 | Left Putamen |
| 7,192 | 0.099 | 4 | 6 | 52 | Right Medial Frontal Gyrus |
| | 0.032 | -2 | 0 | 68 | Left Superior Frontal Gyrus |
| 3,648 | 0.075 | -38 | -16 | 52 | Left Precentral Gyrus |
| 3,192 | 0.092 | 34 | 24 | -2 | Right Insula |
| 2,264 | 0.066 | -30 | 24 | 4 | Left Insula |
| 1,920 | 0.040 | 14 | -86 | -4 | Right Lingual Gyrus |
| | 0.033 | 32 | -86 | -6 | Right Middle Occipital Gyrus |
| | 0.031 | 24 | -90 | -4 | Right Middle Occipital Gyrus |
| 1,696 | 0.051 | 44 | -2 | 48 | Right Precentral Gyrus |
| | 0.026 | 34 | -4 | 52 | Right Precentral Gyrus |
| 1,232 | 0.030 | -28 | -92 | 4 | Left Middle Occipital Gyrus |
| | 0.030 | -24 | -94 | 2 | Left Lingual Gyrus |
| | 0.027 | -18 | -94 | -6 | Left Inferior Occipital Gyrus |
| | 0.024 | -12 | -94 | 2 | Left Lingual Gyrus |
| 1,024 | 0.051 | 36 | 46 | 26 | Right Middle Frontal Gyrus |
| 776 | 0.042 | -36 | 44 | 22 | Left Middle Frontal Gyrus |
| | 0.024 | -32 | 38 | 30 | Left Middle Frontal Gyrus |
| ***Loss anticipation*** | | | | | |
| 25,328 | 0.073 | 12 | 10 | -4 | Right Caudate Head |
| | 0.072 | -10 | 8 | -2 | Left Caudate Head |
| | 0.037 | -8 | -18 | 0 | Left Thalamus |
| | 0.035 | 12 | -2 | 16 | Right Caudate Body |
| | 0.028 | -30 | 22 | -4 | Left Claustrum |
| | 0.022 | 4 | -12 | 8 | Right Thalamus, Medial Dorsal Nucleus |
| | 0.020 | 0 | -6 | 0 | Left Thalamus |
| | 0.018 | -40 | 14 | -2 | Left Insula |
| | 0.017 | 6 | -18 | -2 | Right Thalamus |
| 4,672 | 0.057 | 4 | 2 | 54 | Right Medial Frontal Gyrus |
| | 0.025 | 8 | 16 | 40 | Right Cingulate Gyrus |
| 1,976 | 0.038 | 34 | 24 | 2 | Right Insula |
| | 0.020 | 40 | 14 | 6 | Right Insula |
| 1,712 | 0.034 | -40 | -14 | 56 | Left Precentral Gyrus |
| 1,176 | 0.031 | 42 | -2 | 50 | Right Precentral Gyrus |
| ***Win outcome*** | | | | | |
| 3,280 | 0.035 | 0 | 46 | -4 | Right Anterior Cingulate Cortex |
| 3,160 | 0.036 | -14 | 10 | -12 | Left Putamen |
| | 0.021 | -18 | -6 | -22 | Left Amygdala |
| | 0.018 | -20 | 12 | -4 | Left Putamen |
| 3,072 | 0.047 | 12 | 10 | -10 | Right Caudate Head |
| 2,032 | 0.041 | 2 | -38 | 36 | Left Cingulate Gyrus |
| 1,936 | 0.037 | 26 | -90 | -2 | Right Middle Occipital Gyrus |
| 1,168 | 0.024 | 36 | -64 | 42 | Right Precuneus |
| | 0.021 | 32 | -66 | 34 | Right Precuneus |

| | | | | | |
|---|---|---|---|---|---|
| 1,112 | 0.020 | -16 | -88 | -12 | Left Lingual Gyrus |
| | 0.020 | -28 | -84 | -4 | Left Middle Occipital Gyrus |
| | 0.017 | -24 | -92 | -6 | Left Inferior Occipital Gyrus |
| 992 | 0.043 | 48 | 36 | 16 | Right Middle Frontal Gyrus |
| 824 | 0.025 | -20 | 30 | 48 | Left Superior Frontal Gyrus |
| 656 | 0.024 | -30 | -64 | 40 | Left Precuneus |
| ***Loss outcome*** | | | | | |
| 1,856 | 0.025 | 32 | 20 | -12 | Right Insula |
| | 0.015 | 44 | 22 | -2 | Right Insula |
| | 0.015 | 40 | 22 | -18 | Right Inferior Frontal Gyrus |
| | 0.012 | 42 | 16 | -10 | Right Insula |
| 696 | 0.014 | 0 | 36 | 24 | Left Cingulate Gyrus |
| 584 | 0.016 | -2 | -30 | -4 | Left Thalamus, Pulvinar |

**Supplementary Table S3.** ALE results for conjunction analyses

| Cluster volumes ($mm^3$) | ALE | MNI Coordinates (mm) | | | Identified Region |
|---|---|---|---|---|---|
| | | x | y | z | |
| ***Win & Loss anticipation*** | | | | | |
| 18,336 | 0.073 | 12 | 10 | -4 | Right Caudate Head |
| | 0.072 | -10 | 8 | -2 | Left Caudate Head |
| | 0.035 | -6 | -18 | 2 | Left Thalamus, Medial Dorsal Nucleus |
| | 0.022 | 4 | -12 | 8 | Right Thalamus, Medial Dorsal Nucleus |
| | 0.020 | 0 | -6 | 0 | Left Thalamus |
| | 0.017 | 6 | -18 | -2 | Right Thalamus |
| 3,568 | 0.057 | 4 | 2 | 54 | Right Medial Frontal Gyrus |
| | 0.024 | 8 | 14 | 40 | Right Cingulate Gyrus |
| 1,600 | 0.038 | 34 | 24 | 2 | Right Insula |
| 1,560 | 0.034 | -40 | -14 | 56 | Left Precentral Gyrus |
| 856 | 0.031 | 42 | -2 | 50 | Right Precentral Gyrus |
| 728 | 0.028 | -30 | 22 | -4 | Left Claustrum |
| 8 | 0.021 | -14 | -2 | 14 | Left Caudate Body |
| 8 | 0.015 | 10 | 10 | 38 | Right Cingulate Gyrus |
| ***Win & Loss outcome*** | | | | | |
| NA | | | | | |
| ***Win anticipation & outcome*** | | | | | |
| 3,000 | 0.047 | 12 | 10 | -10 | Right Caudate Head |
| 2,672 | 0.036 | -14 | 10 | -12 | Left Putamen |
| | 0.018 | -20 | 12 | -4 | Left Putamen |
| | 0.016 | -20 | 0 | -18 | Left Parahippocampal Gyrus |
| 872 | 0.031 | 24 | -90 | -4 | Right Middle Occipital Gyrus |
| 24 | 0.016 | -26 | -90 | 0 | Left Middle Occipital Gyrus |
| 8 | 0.014 | -22 | -94 | -8 | Left Inferior Occipital Gyrus |
| 8 | 0.014 | -20 | -92 | -8 | Left Lingual Gyrus |
| 8 | 0.015 | -28 | -88 | 2 | Left Middle Occipital Gyrus |

| | | | | | |
|---|---|---|---|---|---|
| 8 | 0.015 | -26 | -88 | 4 | Left Middle Occipital Gyrus |
| ***Loss anticipation & outcome*** | | | | | |
| 16 | 0.011 | 32 | 22 | -8 | Right Claustrum |
| 16 | 0.010 | 40 | 22 | -2 | Right Insula |

*Note:* NA: not available

**Supplementary Table S4.** ALE results for subtraction analyses

| Cluster volumes (mm³) | Z | MNI Coordinates (mm) | | | Identified Region |
|---|---|---|---|---|---|
| | | x | y | z | |
| ***Win > Loss anticipation*** | | | | | |
| NA | | | | | |
| ***Loss > Win anticipation*** | | | | | |
| NA | | | | | |
| ***Win > Loss outcome*** | | | | | |
| 1,968 | 3.719 | -3.3 | 47.2 | -9.7 | Left Anterior Cingulate Cortex |
| | 3.540 | -1 | 54 | -4 | Left Anterior Cingulate Cortex |
| | 3.090 | 1 | 39 | -12 | Left Anterior Cingulate Cortex |
| ***Loss > Win outcome*** | | | | | |
| NA | | | | | |
| ***Win anticipation > outcome*** | | | | | |
| 6,728 | 3.719 | 1.8 | 6.4 | 1.8 | Left Caudate Head |
| | 0.000 | -2.7 | -13.5 | 4.4 | Left Thalamus, Medial Dorsal Nucleus |
| | 3.353 | 0 | -24 | -2 | Left Midbrain, Red Nucleus |
| | 3.239 | -8 | -12 | 2 | Left Thalamus, Ventral Lateral Nucleus |
| 6,200 | 3.719 | 0.6 | 3.4 | 52.8 | Left Medial Frontal Gyrus |
| 3,128 | 3.719 | -39.5 | -11.7 | 52.4 | Left Precentral Gyrus |
| 1,600 | 3.719 | 45.7 | -2 | 48.3 | Right Precentral Gyrus |
| | 3.540 | 36.5 | -4 | 51 | Right Precentral Gyrus |
| | 3.353 | 38 | -6 | 44 | Right Middle Frontal Gyrus |
| 824 | 3.719 | 36.2 | 29.8 | -1.8 | Right Inferior Frontal Gyrus |
| | 3.540 | 33.3 | 27.1 | -0.8 | Right Insula |
| 304 | 3.719 | 23.8 | 4.8 | -13.7 | Right Putamen |
| 296 | 3.719 | -32 | 16 | 9 | Left Insula |
| | 3.540 | -31 | 17 | 5 | Left Claustrum |
| ***Win outcome > anticipation*** | | | | | |
| 3,152 | 3.719 | -1 | 45.9 | -5.8 | Left Anterior Cingulate Cortex |
| 1,864 | 3.719 | 1 | -37.3 | 35 | Left Cingulate Gyrus |
| 472 | 3.540 | 37.8 | -65.1 | 37.7 | Right Precuneus |
| | 3.239 | 38 | -61 | 44 | Right Inferior Parietal Lobule |
| 400 | 3.540 | 47 | 36.4 | 14.1 | Right Middle Frontal Gyrus |
| 184 | 3.719 | -17.5 | 32.5 | 46 | Left Superior Frontal Gyrus |
| | 3.156 | -22 | 34 | 44 | Left Superior Frontal Gyrus |
| ***Loss anticipation > outcome*** | | | | | |

| | | | | | |
|---|---|---|---|---|---|
| 4,056 | 3.719 | -15.5 | 11.3 | -9.1 | Left Putamen |
| | 3.239 | -15.5 | 4 | -7 | Left Lateral Globus Pallidus |
| | 3.353 | -8 | 15 | -4 | Left Caudate Head |
| 3,440 | 3.719 | 16.3 | 10.2 | -9.3 | Right Putamen |
| ***Loss outcome > anticipation*** | | | | | |
| NA | | | | | |

*Note:* NA: not available

**Supplementary Table S5.** ALE results for activations during win anticipation, loss anticipation, win outcome, and loss outcome, for the evaluation of publication bias

| Cluster volumes ($mm^3$) | ALE | MNI Coordinates (mm) | | | Identified Region |
|---|---|---|---|---|---|
| | | x | y | z | |
| ***Win anticipation*** | | | | | |
| 14,688 | 0.20 | 10 | 10 | -2 | Right Caudate Head |
| | 0.16 | -10 | 10 | -4 | Left Caudate Head |
| | 0.07 | 4 | -10 | 6 | Right Thalamus, Medial Dorsal Nucleus |
| | 0.07 | -10 | -16 | 8 | Left Thalamus, Medial Dorsal Nucleus |
| | 0.07 | -18 | -4 | 12 | Left Thalamus, Ventral Anterior Nucleus |
| | 0.07 | -6 | -8 | 8 | Left Thalamus, Anterior Nucleus |
| 3,992 | 0.12 | 4 | 6 | 52 | Right Medial Frontal Gyrus |
| ***Loss anticipation*** | | | | | |
| 13,504 | 0.09 | 12 | 10 | -2 | Right Caudate Head |
| | 0.08 | -10 | 8 | -2 | Left Caudate Head |
| | 0.05 | -10 | -2 | 12 | Left Caudate Body |
| | 0.05 | 12 | -2 | 14 | Right Caudate Body |
| | 0.04 | 4 | -12 | 10 | Right Thalamus |
| | 0.04 | -8 | -16 | 0 | Left Thalamus |
| | 0.04 | 24 | 4 | 4 | Right Putamen |
| | 0.04 | 20 | 2 | -12 | Right Lateral Globus Pallidus |
| ***Win outcome*** | | | | | |
| Nil | | | | | |
| ***Loss outcome*** | | | | | |
| 728 | 0.03 | 32 | 20 | -12 | Right Insula |
| | 0.02 | 40 | 20 | -14 | Right Inferior Frontal Gyrus |

*Note:* Nil: no significant findings

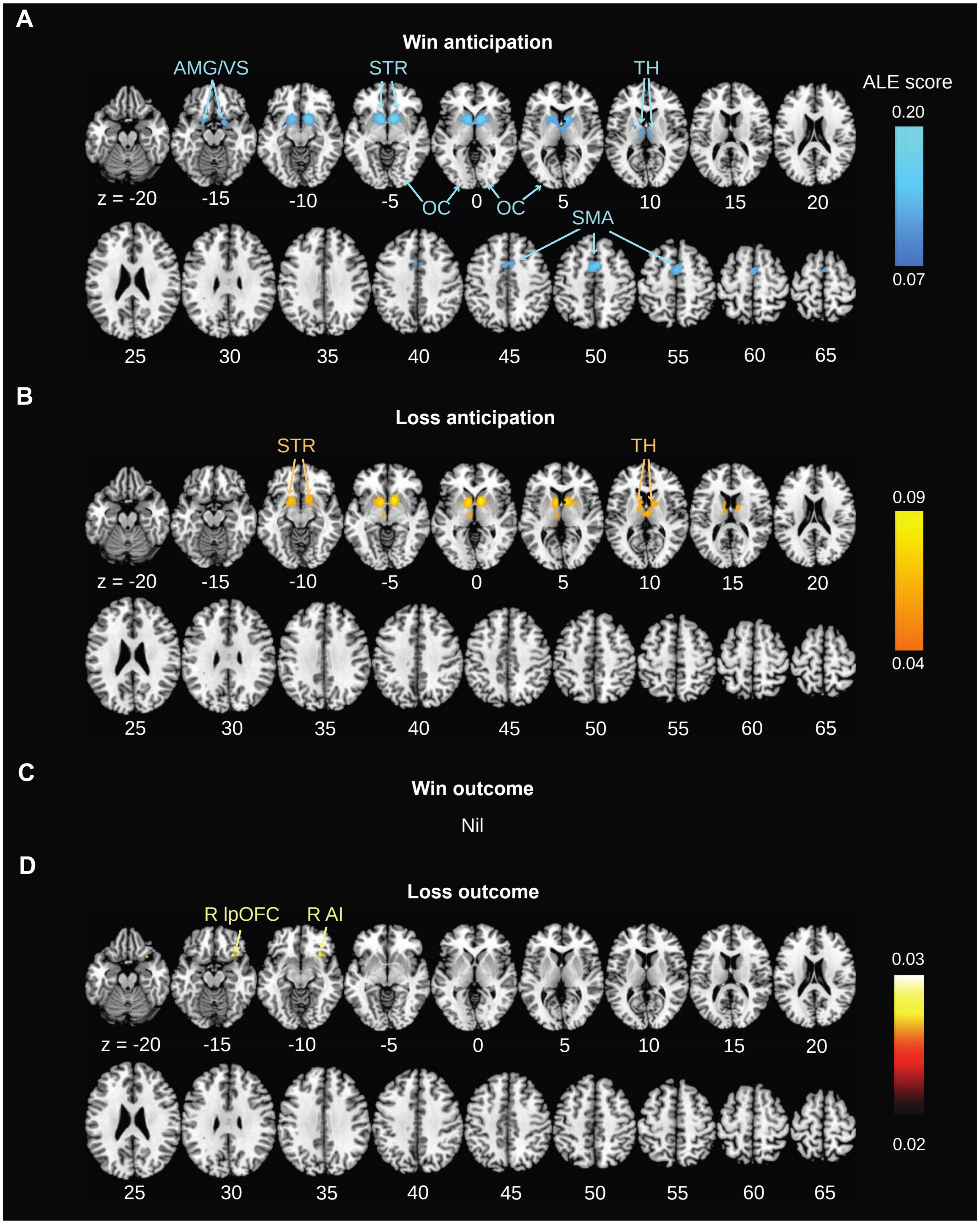


**Supplementary Figure S1.** ALE single dataset analyses for publication bias evaluation. **(A)** Win anticipation; **(B)** Loss anticipation; **(C)** Win outcome; and **(D)** Loss outcome. *Note:* The results were evaluated with a cluster-forming threshold of $p < 0.001$ uncorrected and a cluster-level threshold of $p < 0.05$ FWE corrected. Color bars represent ALE scores. Nil: no significant findings; R: right; AI: anterior insula; AMG: amygdala; lpOFC: lateral posterior orbitofrontal cortex; OC: occipital cortex; SMA: supplementary motor area; STR: striatum; TH: thalamus; VS: ventral striatum

### References

Abler, B., Erk, S., Walter, H., 2007. Human reward system activation is modulated by a single dose of olanzapine in healthy subjects in an event-related, double-blind, placebo-controlled fMRI study. Psychopharmacology (Berl) 191, 823-833.

Adcock, R.A., Thangavel, A., Whitfield-Gabrieli, S., Knutson, B., Gabrieli, J.D., 2006. Reward-motivated learning: mesolimbic activation precedes memory formation. Neuron 50, 507-517.

Asari, Y., Ikeda, Y., Tateno, A., Okubo, Y., Iijima, T., Suzuki, H., 2018. Acute tramadol enhances brain activity associated with reward anticipation in the nucleus accumbens. Psychopharmacology (Berl) 235, 2631-2642.

Balodis, I.M., Kober, H., Worhunsky, P.D., Stevens, M.C., Pearlson, G.D., Potenza, M.N., 2012. Diminished frontostriatal activity during processing of monetary rewards and losses in pathological gambling. Biol Psychiatry 71, 749-757.

Barker, E.D., Biondo, F., Jia, T., Pingault, J.-B., Du Rietz, E., Zhang, Y., Ruggeri, B., Banaschewski, T., Hohmann, S., Bokde, A.L., 2021. Do ADHD-impulsivity and BMI have shared polygenic and neural correlates? Molecular psychiatry 26, 1019-1028.

Barman, A., Richter, S., Soch, J., Deibele, A., Richter, A., Assmann, A., Wustenberg, T., Walter, H., Seidenbecher, C.I., Schott, B.H., 2015. Gender-specific modulation of neural mechanisms underlying social reward processing by Autism Quotient. Soc Cogn Affect Neurosci 10, 1537-1547.

Behan, B., Stone, A., Garavan, H., 2015. Right prefrontal and ventral striatum interactions underlying impulsive choice and impulsive responding. Hum Brain Mapp 36, 187-198.

Bhutani, S., Christian, I.R., Palumbo, D., Wiggins, J.L., 2021. Reward-related neural correlates in adolescents with excess body weight. Neuroimage Clin 30, 102618.

Bjork, J.M., Smith, A.R., Chen, G., Hommer, D.W., 2010. Adolescents, adults and rewards: comparing motivational neurocircuitry recruitment using fMRI. PLoS One 5, e11440.

Bretzke, M., Wahl, H., Plichta, M.M., Wolff, N., Roessner, V., Vetter, N.C., Buse, J., 2021. Ventral Striatal Activation During Reward Anticipation of Different Reward Probabilities in Adolescents and Adults. Front Hum Neurosci 15, 649724.

Cao, Z., Bennett, M., Orr, C., Icke, I., Banaschewski, T., Barker, G.J., Bokde, A.L.W., Bromberg, U., Buchel, C., Quinlan, E.B., Desrivieres, S., Flor, H., Frouin, V., Garavan, H., Gowland, P., Heinz, A., Ittermann, B., Martinot, J.L., Nees, F., Orfanos, D.P., Paus, T., Poustka, L., Hohmann, S., Frohner, J.H., Smolka, M.N., Walter, H., Schumann, G., Whelan, R., Consortium, I., 2019. Mapping adolescent reward anticipation, receipt, and prediction error during the monetary incentive delay task. Hum Brain Mapp 40, 262-283.

Carl, H., Walsh, E., Eisenlohr-Moul, T., Minkel, J., Crowther, A., Moore, T., Gibbs, D., Petty, C., Bizzell, J., Dichter, G.S., Smoski, M.J., 2016. Sustained anterior cingulate cortex activation during reward processing predicts response to psychotherapy in major depressive disorder. J Affect Disord 203, 204-212.

Carter, R.M., Macinnes, J.J., Huettel, S.A., Adcock, R.A., 2009. Activation in the VTA and nucleus accumbens increases in anticipation of both gains and losses. Front Behav Neurosci 3, 21.

Cho, Y.T., Fromm, S., Guyer, A.E., Detloff, A., Pine, D.S., Fudge, J.L., Ernst, M., 2013. Nucleus accumbens, thalamus and insula connectivity during incentive anticipation in typical adults and adolescents. Neuroimage 66, 508-521.

Cohn, M.D., Veltman, D.J., Pape, L.E., van Lith, K., Vermeiren, R.R., van den Brink, W., Doreleijers, T.A., Popma, A., 2015. Incentive Processing in Persistent Disruptive Behavior and Psychopathic Traits: A Functional Magnetic Resonance Imaging Study in Adolescents. Biol Psychiatry 78, 615-624.

Colich, N.L., Ho, T.C., Ellwood-Lowe, M.E., Foland-Ross, L.C., Sacchet, M.D., LeMoult, J.L., Gotlib, I.H., 2017. Like mother like daughter: putamen activation as a mechanism underlying intergenerational risk for depression. Soc Cogn Affect Neurosci 12, 1480-1489.

Cope, L.M., Martz, M.E., Hardee, J.E., Zucker, R.A., Heitzeg, M.M., 2019. Reward activation in childhood predicts adolescent substance use initiation in a high-risk sample. Drug Alcohol Depend 194, 318-325.

Costumero, V., Barros-Loscertales, A., Bustamante, J.C., Ventura-Campos, N., Fuentes, P., Avila, C., 2013. Reward sensitivity modulates connectivity among reward brain areas during processing of anticipatory reward cues. Eur J Neurosci 38, 2399-2407.

Damiano, C.R., Aloi, J., Dunlap, K., Burrus, C.J., Mosner, M.G., Kozink, R.V., McLaurin, R.E., Mullette-Gillman, O., Carter, R.M., Huettel, S.A., 2014. Association between the oxytocin receptor (OXTR) gene and mesolimbic responses to rewards. Molecular autism 5, 1-12.
Dillon, D.G., Bogdan, R., Fagerness, J., Holmes, A.J., Perlis, R.H., Pizzagalli, D.A., 2010. Variation in TREK1 gene linked to depression-resistant phenotype is associated with potentiated neural responses to rewards in humans. Hum Brain Mapp 31, 210-221.
Dunlop, K., Rizvi, S.J., Kennedy, S.H., Hassel, S., Strother, S.C., Harris, J.K., Zamyadi, M., Arnott, S.R., Davis, A.D., Mansouri, F., Schulze, L., Ceniti, A.K., Lam, R.W., Milev, R., Rotzinger, S., Foster, J.A., Frey, B.N., Parikh, S.V., Soares, C.N., Uher, R., Turecki, G., MacQueen, G.M., Downar, J., 2020. Clinical, behavioral, and neural measures of reward processing correlate with escitalopram response in depression: a Canadian Biomarker Integration Network in Depression (CAN-BIND-1) Report. Neuropsychopharmacology 45, 1390-1397.
Edel, M.A., Enzi, B., Witthaus, H., Tegenthoff, M., Peters, S., Juckel, G., Lissek, S., 2013. Differential reward processing in subtypes of adult attention deficit hyperactivity disorder. J Psychiatr Res 47, 350-356.
Enzi, B., Edel, M.A., Lissek, S., Peters, S., Hoffmann, R., Nicolas, V., Tegenthoff, M., Juckel, G., Saft, C., 2012. Altered ventral striatal activation during reward and punishment processing in premanifest Huntington's disease: a functional magnetic resonance study. Exp Neurol 235, 256-264.
Figee, M., Vink, M., de Geus, F., Vulink, N., Veltman, D.J., Westenberg, H., Denys, D., 2011. Dysfunctional reward circuitry in obsessive-compulsive disorder. Biol Psychiatry 69, 867-874.
Filbey, F.M., Dunlop, J., Myers, U.S., 2013. Neural effects of positive and negative incentives during marijuana withdrawal. PLoS One 8, e61470.
Funayama, T., Ikeda, Y., Tateno, A., Takahashi, H., Okubo, Y., Fukayama, H., Suzuki, H., 2014. Modafinil augments brain activation associated with reward anticipation in the nucleus accumbens. Psychopharmacology (Berl) 231, 3217-3228.

Gonzalez, M.Z., Allen, J.P., Coan, J.A., 2016. Lower neighborhood quality in adolescence predicts higher mesolimbic sensitivity to reward anticipation in adulthood. Developmental cognitive neuroscience 22, 48-57.
Grimm, O., Nagele, M., Kupper-Tetzel, L., de Greck, M., Plichta, M., Reif, A., 2021. No effect of a dopaminergic modulation fMRI task by amisulpride and L-DOPA on reward anticipation in healthy volunteers. Psychopharmacology (Berl) 238, 1333-1342.
Hanssen, E., van der Velde, J., Gromann, P.M., Shergill, S.S., de Haan, L., Bruggeman, R., Krabbendam, L., Aleman, A., van Atteveldt, N., 2015. Neural correlates of reward processing in healthy siblings of patients with schizophrenia. Front Hum Neurosci 9, 504.
He, Z., Zhang, D., Muhlert, N., Elliott, R., 2019. Neural substrates for anticipation and consumption of social and monetary incentives in depression. Soc Cogn Affect Neurosci 14, 815-826.
Herbort, M.C., Soch, J., Wustenberg, T., Krauel, K., Pujara, M., Koenigs, M., Gallinat, J., Walter, H., Roepke, S., Schott, B.H., 2016. A negative relationship between ventral striatal loss anticipation response and impulsivity in borderline personality disorder. Neuroimage Clin 12, 724-736.
Ikeda, Y., Funayama, T., Tateno, A., Fukayama, H., Okubo, Y., Suzuki, H., 2019. Bupropion increases activation in nucleus accumbens during anticipation of monetary reward. Psychopharmacology (Berl) 236, 3655-3665.
Johnson, S.L., Mehta, H., Ketter, T.A., Gotlib, I.H., Knutson, B., 2019. Neural responses to monetary incentives in bipolar disorder. Neuroimage Clin 24, 102018.
Juckel, G., Friedel, E., Koslowski, M., Witthaus, H., Ozgurdal, S., Gudlowski, Y., Knutson, B., Wrase, J., Brune, M., Heinz, A., Schlagenhauf, F., 2012. Ventral striatal activation during reward processing in subjects with ultra-high risk for schizophrenia. Neuropsychobiology 66, 50-56.
Juckel, G., Schlagenhauf, F., Koslowski, M., Wustenberg, T., Villringer, A., Knutson, B., Wrase, J., Heinz, A., 2006. Dysfunction of ventral striatal reward prediction in schizophrenia. Neuroimage 29, 409-416.

Kappel, V., Koch, A., Lorenz, R.C., Bruhl, R., Renneberg, B., Lehmkuhl, U., Salbach-Andrae, H., Beck, A., 2013. CID: a valid incentive delay paradigm for children. J Neural Transm (Vienna) 120, 1259-1270.
Kaufmann, C., Beucke, J.C., Preusse, F., Endrass, T., Schlagenhauf, F., Heinz, A., Juckel, G., Kathmann, N., 2013. Medial prefrontal brain activation to anticipated reward and loss in obsessive-compulsive disorder. Neuroimage Clin 2, 212-220.
Kim, M., Mawla, I., Albrecht, D.S., Admon, R., Torrado-Carvajal, A., Bergan, C., Protsenko, E., Kumar, P., Edwards, R.R., Saha, A., Napadow, V., Pizzagalli, D.A., Loggia, M.L., 2020. Striatal hypofunction as a neural correlate of mood alterations in chronic pain patients. Neuroimage 211, 116656.
Kirk, U., Brown, K.W., Downar, J., 2015. Adaptive neural reward processing during anticipation and receipt of monetary rewards in mindfulness meditators. Soc Cogn Affect Neurosci 10, 752-759.
Knutson, B., Bhanji, J.P., Cooney, R.E., Atlas, L.Y., Gotlib, I.H., 2008. Neural responses to monetary incentives in major depression. Biological Psychiatry 63, 686-692.
Kocsel, N., Galambos, A., Szabo, E., Edes, A.E., Magyar, M., Zsombok, T., Pap, D., Kozak, L.R., Bagdy, G., Kokonyei, G., Juhasz, G., 2019. Altered neural activity to monetary reward/loss processing in episodic migraine. Sci Rep 9, 5420.
Kocsel, N., Szabo, E., Galambos, A., Edes, A., Pap, D., Elliott, R., Kozak, L.R., Bagdy, G., Juhasz, G., Kokonyei, G., 2017. Trait Rumination Influences Neural Correlates of the Anticipation but Not the Consumption Phase of Reward Processing. Front Behav Neurosci 11, 85.
Kollmann, B., Scholz, V., Linke, J., Kirsch, P., Wessa, M., 2017. Reward anticipation revisited- evidence from an fMRI study in euthymic bipolar I patients and healthy first-degree relatives. J Affect Disord 219, 178-186.
Li, Z., Wang, Y., Yan, C., Cheung, E.F., Docherty, A.R., Sham, P.C., Gur, R.E., Gur, R.C., Chan, R.C., 2019. Inheritance of neural substrates for motivation and pleasure. Psychological science 30, 1205-1217.
Maresh, E.L., Allen, J.P., Coan, J.A., 2014. Increased default mode network activity in socially anxious individuals during reward processing. Biology of mood & anxiety disorders 4, 1-12.

Martz, M.E., Cope, L.M., Hardee, J.E., Brislin, S.J., Weigard, A., Zucker, R.A., Heitzeg, M.M., 2021. Subtypes of inhibitory and reward activation associated with substance use variation in adolescence: A latent profile analysis of brain imaging data. Cogn Affect Behav Neurosci 21, 1101-1114.
Martz, M.E., Zucker, R.A., Schulenberg, J.E., Heitzeg, M.M., 2018. Psychosocial and neural indicators of resilience among youth with a family history of substance use disorder. Drug Alcohol Depend 185, 198-206.
Metzak, P.D., Addington, J., Hassel, S., Goldstein, B.I., MacIntosh, B.J., Lebel, C., Wang, J.L., Kennedy, S.H., MacQueen, G.M., Bray, S., 2021. Functional imaging in youth at risk for transdiagnostic serious mental illness: Initial results from the PROCAN study. Early Interv Psychiatry 15, 1276-1291.
Michielse, S., Lange, I., Bakker, J., Goossens, L., Verhagen, S., Papalini, S., Wichers, M., Lieverse, R., Schruers, K., van Amelsvoort, T., van Os, J., Murray, G.K., Marcelis, M., 2019. Reward anticipation in individuals with subclinical psychotic experiences: A functional MRI approach. Eur Neuropsychopharmacol 29, 1374-1385.
Montoya, E.R., Bos, P.A., Terburg, D., Rosenberger, L.A., van Honk, J., 2014. Cortisol administration induces global down-regulation of the brain's reward circuitry. Psychoneuroendocrinology 47, 31-42.
Morelli, N.M., Liuzzi, M.T., Duong, J.B., Kryza-Lacombe, M., Chad-Friedman, E., Villodas, M.T., Dougherty, L.R., Wiggins, J.L., 2021. Reward-related neural correlates of early life stress in school-aged children. Dev Cogn Neurosci 49, 100963.
Mucci, A., Dima, D., Soricelli, A., Volpe, U., Bucci, P., Frangou, S., Prinster, A., Salvatore, M., Galderisi, S., Maj, M., 2015. Is avolition in schizophrenia associated with a deficit of dorsal caudate activity? A functional magnetic resonance imaging study during reward anticipation and feedback. Psychol Med 45, 1765-1778.
Murray, L., Lopez-Duran, N.L., Mitchell, C., Monk, C.S., Hyde, L.W., 2020. Neural mechanisms of reward and loss processing in a low-income sample of at-risk adolescents. Soc Cogn Affect Neurosci 15, 1310-1325.
Navas, J.F., Barros-Loscertales, A., Costumero-Ramos, V., Verdejo-Roman, J., Vilar-Lopez, R., Verdejo-Garcia, A., 2018. Excessive body fat linked to blunted

somatosensory cortex response to general reward in adolescents. Int J Obes (Lond) 42, 88-94.
Nymberg, C., Jia, T., Lubbe, S., Ruggeri, B., Desrivieres, S., Barker, G., Buchel, C., Fauth-Buehler, M., Cattrell, A., Conrod, P., Flor, H., Gallinat, J., Garavan, H., Heinz, A., Ittermann, B., Lawrence, C., Mann, K., Nees, F., Salatino-Oliveira, A., Paillere Martinot, M.L., Paus, T., Rietschel, M., Robbins, T., Smolka, M., Banaschewski, T., Rubia, K., Loth, E., Schumann, G., Consortium, I., 2013. Neural mechanisms of attention-deficit/hyperactivity disorder symptoms are stratified by MAOA genotype. Biol Psychiatry 74, 607-614.
Ossewaarde, L., van Wingen, G.A., Kooijman, S.C., Backstrom, T., Fernandez, G., Hermans, E.J., 2011. Changes in functioning of mesolimbic incentive processing circuits during the premenstrual phase. Soc Cogn Affect Neurosci 6, 612-620.
Paraskevopoulou, M., van Rooij, D., Batalla, A., Chauvin, R., Luijten, M., Schene, A.H., Buitelaar, J.K., Schellekens, A.F.A., 2021. Effects of substance misuse on reward-processing in patients with attention-deficit/hyperactivity disorder. Neuropsychopharmacology 46, 622-631.
Pfabigan, D.M., Seidel, E.M., Sladky, R., Hahn, A., Paul, K., Grahl, A., Kublbock, M., Kraus, C., Hummer, A., Kranz, G.S., Windischberger, C., Lanzenberger, R., Lamm, C., 2014. P300 amplitude variation is related to ventral striatum BOLD response during gain and loss anticipation: an EEG and fMRI experiment. Neuroimage 96, 12-21.
Rademacher, L., Krach, S., Kohls, G., Irmak, A., Grunder, G., Spreckelmeyer, K.N., 2010. Dissociation of neural networks for anticipation and consumption of monetary and social rewards. Neuroimage 49, 3276-3285.
Saji, K., Ikeda, Y., Kim, W., Shingai, Y., Tateno, A., Takahashi, H., Okubo, Y., Fukayama, H., Suzuki, H., 2013. Acute NK(1) receptor antagonist administration affects reward incentive anticipation processing in healthy volunteers. Int J Neuropsychopharmacol 16, 1461-1471.
Schlagenhauf, F., Juckel, G., Koslowski, M., Kahnt, T., Knutson, B., Dembler, T., Kienast, T., Gallinat, J., Wrase, J., Heinz, A., 2008. Reward system activation in schizophrenic patients switched from typical neuroleptics to olanzapine. Psychopharmacology (Berl) 196, 673-684.

Simon, J.J., Skunde, M., Wu, M., Schnell, K., Herpertz, S.C., Bendszus, M., Herzog, W., Friederich, H.C., 2015. Neural dissociation of food- and money-related reward processing using an abstract incentive delay task. Soc Cogn Affect Neurosci 10, 1113-1120.

Simon, J.J., Walther, S., Fiebach, C.J., Friederich, H.C., Stippich, C., Weisbrod, M., Kaiser, S., 2010. Neural reward processing is modulated by approach- and avoidance-related personality traits. Neuroimage 49, 1868-1874.

Stoy, M., Schlagenhauf, F., Schlochtermeier, L., Wrase, J., Knutson, B., Lehmkuhl, U., Huss, M., Heinz, A., Strohle, A., 2011. Reward processing in male adults with childhood ADHD--a comparison between drug-naive and methylphenidate-treated subjects. Psychopharmacology (Berl) 215, 467-481.

Stoy, M., Schlagenhauf, F., Sterzer, P., Bermpohl, F., Hagele, C., Suchotzki, K., Schmack, K., Wrase, J., Ricken, R., Knutson, B., Adli, M., Bauer, M., Heinz, A., Strohle, A., 2012. Hyporeactivity of ventral striatum towards incentive stimuli in unmedicated depressed patients normalizes after treatment with escitalopram. J Psychopharmacol 26, 677-688.

Strohle, A., Stoy, M., Wrase, J., Schwarzer, S., Schlagenhauf, F., Huss, M., Hein, J., Nedderhut, A., Neumann, B., Gregor, A., Juckel, G., Knutson, B., Lehmkuhl, U., Bauer, M., Heinz, A., 2008. Reward anticipation and outcomes in adult males with attention-deficit/hyperactivity disorder. Neuroimage 39, 966-972.

Treadway, M.T., Buckholtz, J.W., Zald, D.H., 2013. Perceived stress predicts altered reward and loss feedback processing in medial prefrontal cortex. Front Hum Neurosci 7, 180.

van Hell, H.H., Jager, G., Bossong, M.G., Brouwer, A., Jansma, J.M., Zuurman, L., van Gerven, J., Kahn, R.S., Ramsey, N.F., 2012. Involvement of the endocannabinoid system in reward processing in the human brain. Psychopharmacology (Berl) 219, 981-990.

Verdejo-Roman, J., Fornito, A., Soriano-Mas, C., Vilar-Lopez, R., Verdejo-Garcia, A., 2017. Independent functional connectivity networks underpin food and monetary reward sensitivity in excess weight. Neuroimage 146, 293-300.

Veroude, K., von Rhein, D., Chauvin, R.J., van Dongen, E.V., Mennes, M.J., Franke, B., Heslenfeld, D.J., Oosterlaan, J., Hartman, C.A., Hoekstra, P.J., Glennon, J.C., Buitelaar, J.K., 2016. The link between callous-unemotional traits and neural mechanisms of reward processing: An fMRI study. Psychiatry Res Neuroimaging 255, 75-80.

Weidacker, K., Kim, S.G., Nord, C.L., Rua, C., Rodgers, C.T., Voon, V., 2021. Avoiding monetary loss: A human habenula functional MRI ultra-high field study. Cortex 142, 62-73.

Weiland, B.J., Heitzeg, M.M., Zald, D., Cummiford, C., Love, T., Zucker, R.A., Zubieta, J.K., 2014. Relationship between impulsivity, prefrontal anticipatory activation, and striatal dopamine release during rewarded task performance. Psychiatry Res 223, 244-252.

Wu, C.C., Samanez-Larkin, G.R., Katovich, K., Knutson, B., 2014. Affective traits link to reliable neural markers of incentive anticipation. Neuroimage 84, 279-289.

Yan, C., Wang, Y., Su, L., Xu, T., Yin, D.Z., Fan, M.X., Deng, C.P., Wang, Z.X., Lui, S.S., Cheung, E.F., Chan, R.C., 2016. Differential mesolimbic and prefrontal alterations during reward anticipation and consummation in positive and negative schizotypy. Psychiatry Res Neuroimaging 254, 127-136.

Yao, Y.W., Liu, L., Worhunsky, P.D., Lichenstein, S., Ma, S.S., Zhu, L., Shi, X.H., Yang, S., Zhang, J.T., Yip, S.W., 2020. Is monetary reward processing altered in drug-naive youth with a behavioral addiction? Findings from internet gaming disorder. Neuroimage Clin 26, 102202.

Yau, W.Y., Zubieta, J.K., Weiland, B.J., Samudra, P.G., Zucker, R.A., Heitzeg, M.M., 2012. Nucleus accumbens response to incentive stimuli anticipation in children of alcoholics: relationships with precursive behavioral risk and lifetime alcohol use. J Neurosci 32, 2544-2551.

Zweynert, S., Pade, J.P., Wustenberg, T., Sterzer, P., Walter, H., Seidenbecher, C.I., Richardson-Klavehn, A., Duzel, E., Schott, B.H., 2011. Motivational salience modulates hippocampal repetition suppression and functional connectivity in humans. Front Hum Neurosci 5, 144.
